# Whole Animal Multiplexed Single-Cell RNA-Seq Reveals Plasticity of *Clytia* Medusa Cell Types

**DOI:** 10.1101/2021.01.22.427844

**Authors:** Tara Chari, Brandon Weissbourd, Jase Gehring, Anna Ferraioli, Lucas Leclère, Makenna Herl, Fan Gao, Sandra Chevalier, Richard R. Copley, Evelyn Houliston, David J. Anderson, Lior Pachter

## Abstract

We present an organism-wide, transcriptomic cell atlas of the hydrozoan medusa *Clytia hemisphaerica*, and determine how its component cell types respond to starvation. Utilizing multiplexed scRNA-seq, in which individual animals were indexed and pooled from control and perturbation conditions into a single sequencing run, we avoid artifacts from batch effects and are able to discern shifts in cell state in response to organismal perturbations. This work serves as a foundation for future studies of development, function, and plasticity in a genetically tractable jellyfish species. Moreover, we introduce a powerful workflow for high-resolution, <u>wh</u>ole <u>a</u>nimal, <u>m</u>ultiplexed single-cell genomics (WHAM-seq) that is readily adaptable to other traditional or non-traditional model organisms.

## Introduction

Single-cell RNA sequencing (scRNA-seq) is enabling the survey of complete transcriptomes of thousands to millions of cells^1^, resulting in the establishment of cell atlases across whole organisms^2–6^, exploration of the diversity of cell types throughout the animal kingdom^3,7–9^, and investigation of transcriptomic changes under perturbation^10,11^. However, scRNA-seq studies involving multiple samples can be costly, and may be confounded by batch effects resulting from multiple distinct library preparations^12,13^. Recent developments in scRNA-seq multiplexing technology expand the number of samples, individuals, or perturbations that can be incorporated within runs, facilitating well-controlled scRNA-seq experiments^11,14–18^. These advances have created an opportunity to explore systems biology of whole organisms at single-cell resolution, merging the concepts of cell atlas surveys with multiplexed single-cell experimentation.

Here we apply this powerful experimental paradigm to interrogate a planktonic model organism. We examine the medusa (free-swimming jellyfish) stage of the hydrozoan *Clytia hemisphaerica*, with dual motivations. Firstly, *Clytia* is a powerful, emerging model system spanning multiple fields, from evolutionary and developmental biology to regeneration and neuroscience^19–24^. While previous work has characterized a number of cell types in the *Clytia* medusa^21^, a whole-organism atlas of transcriptomic cell types has been lacking. Such an atlas is a critical foundation for the *Clytia* community, and an important addition to the study of cell types across animal phylogeny.

Secondly, emerging multiplexing techniques present new opportunities for systems-level studies of cell types and their changing states at unprecedented resolution in whole organisms. The *Clytia* medusa offers an appealing platform for pioneering such studies. It is small, transparent, and has simple tissues and organs, stem cell populations actively replenishing many cell types in mature animals, and remarkable tissue plasticity and regenerative capacity^19,22,24–27^. Furthermore, the 1cm-diameter adult medusae used in this study contain on the order of 10^5^ cells (see Methods), making it possible to sample cells comprehensively across a whole animal in a cost-effective manner (Supplementary Table 1) using current scRNA-seq technology. In this study, we generate a cell atlas for the *Clytia* medusa while simultaneously performing a whole organism perturbation study, providing the first medusa single-cell dataset and an examination of changing cell states across the organism. Our work also provides a proof-of-principle for perturbation studies in non-traditional model organisms, using multiplexing technology and a reproducible workflow, from the experimental implementation to the data processing and analysis.

## Results

We compared control versus starved animals, as this strong, naturalistic stimulus was likely to cause significant, interpretable changes in transcription across multiple cell types. Laboratory-raised, young adult, female medusae were split into two groups of five animals, one deprived of food for four days, and the second fed daily (see Methods). We observed numerous phenotypic changes in starved animals, including a dramatic size reduction reflecting ∼five-fold fewer total cells^28^ (Fig. 1; see Methods), and a striking reduction in gonad size. Correspondingly, the number of eggs released per day decreased^29^ (Supplementary Fig. 1).

**Figure 1:**
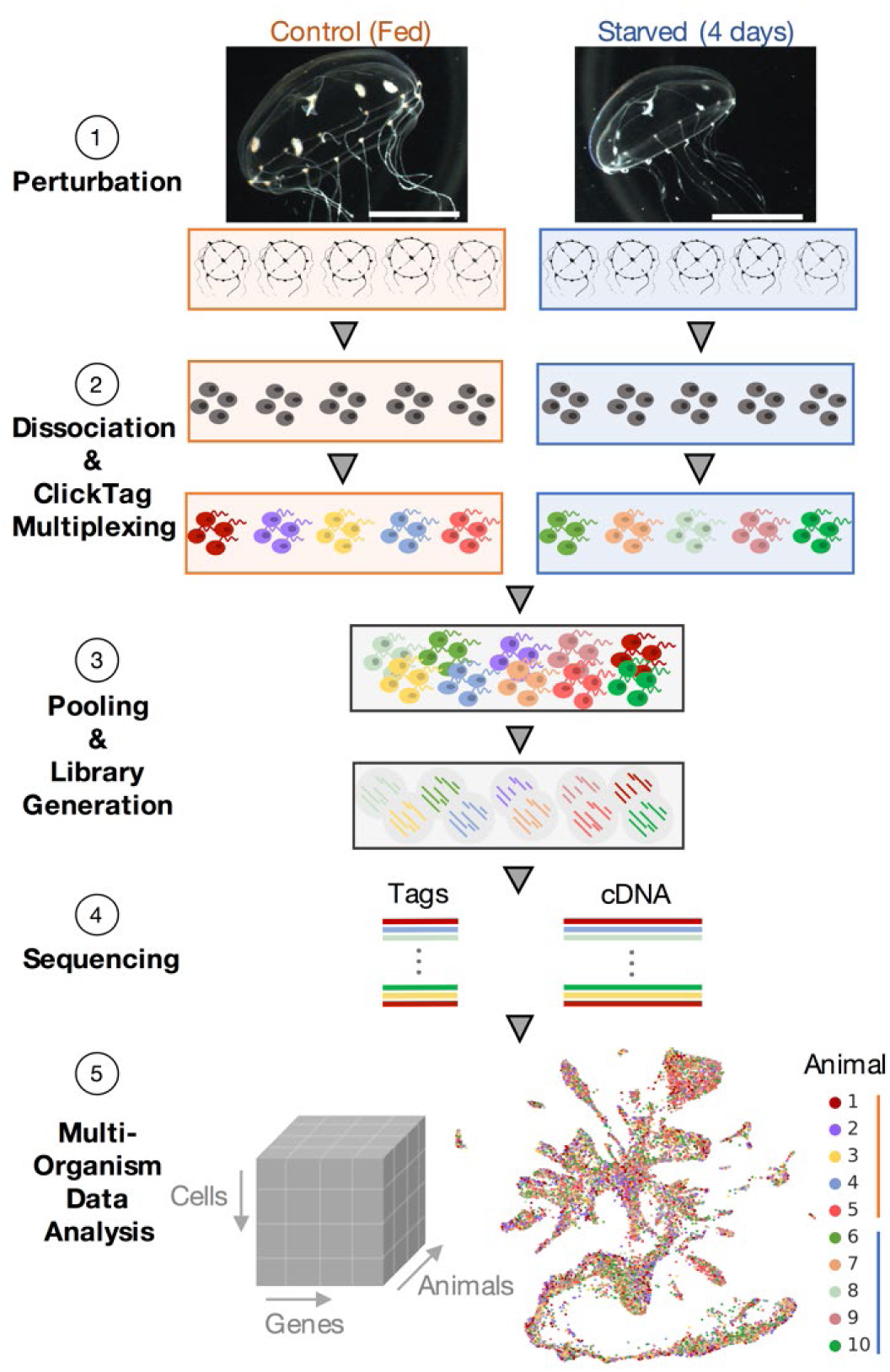
Overview of Whole Organism Multiplexed Experimentation: Experimental design of the starvation experiment showing 1) images of control (fed) versus 4-day starved animals (scale bars represent 0.5 cm), 2) dissociation of individual medusa and chemical tagging of cells with ClickTags to enable multiplexed scRNA-seq, 3) pooling of cells and library generation from lysed cells to generate 4) sequencing libraries for the multiplexed cDNA and ClickTag data and create 5) single-cell resolved gene expression count matrices from all animals (see Methods). [[Code for 5](https://github.com/pachterlab/CWGFLHGCCHAP_2021/blob/master/notebooks/CellAtlasAnalysis/starvation_Analysis.ipynb)]

For scRNA-seq, single-cell suspensions were prepared from each whole medusa and individually labeled with unique ClickTag barcodes^14^ using a sea-water compatible workflow (see Methods; Supplementary Table 2,3; Supplementary Methods). All labeled suspensions were pooled and processed with the 10X Genomics V2.0 workflow and Illumina sequencing, allowing construction of a combined dataset across organisms and treatments, without the necessity of batch correction (Fig. 1; Supplementary Table 1; Supplementary Fig. 2). A total of 13,673 single-cell profiles derived from 10 individuals (5 control, 5 starved) passed quality control, with high concordance in cell type abundance and gene expression among animals in the same treatment condition (see Methods, Supplementary Fig. 3). From this gene expression matrix, we 1) derived a *Clytia* medusa cell atlas, and 2) performed a high-resolution analysis of the transcriptional impact of starvation across all observed cell types. To assess the utility of this approach, we performed a separate experiment examining *Clytia* medusae responses to multiple transient stimuli over short timescales (Supplementary Fig. 4-9, Supplementary Table 4). This enabled the examination of batch effects across and within multiplexed experiments (Supplementary Fig. 10).

### A Clytia Cell Atlas

To generate the cell atlas, we clustered the cells using the gene expression matrix, extracting 36 cell types present across all individuals, along with their corresponding marker genes (Fig. 2a,b; Supplementary Fig. 11,12,14; Supplementary Table 5; see Methods). We then generated a low-dimensional representation^30,31^ of these cell types (Fig. 2a). We could group the cell types into eight broad classes (Fig. 2a) by considering their individual deduced identities (see below) as well as shared marker gene expression. These broad classes correspond to the two cnidarian body layers (the outer epidermis or inner gastrodermis) and likely derivatives of the multipotent interstitial stem cell population (i-cells). I-cells are a specific feature of hydrozoans, and are particularly well characterized in Hydra, where they generate neural cells, gland cells, and stinging cells (nematocytes), as well as germ cells^8,20,32^.

**Figure 2:**
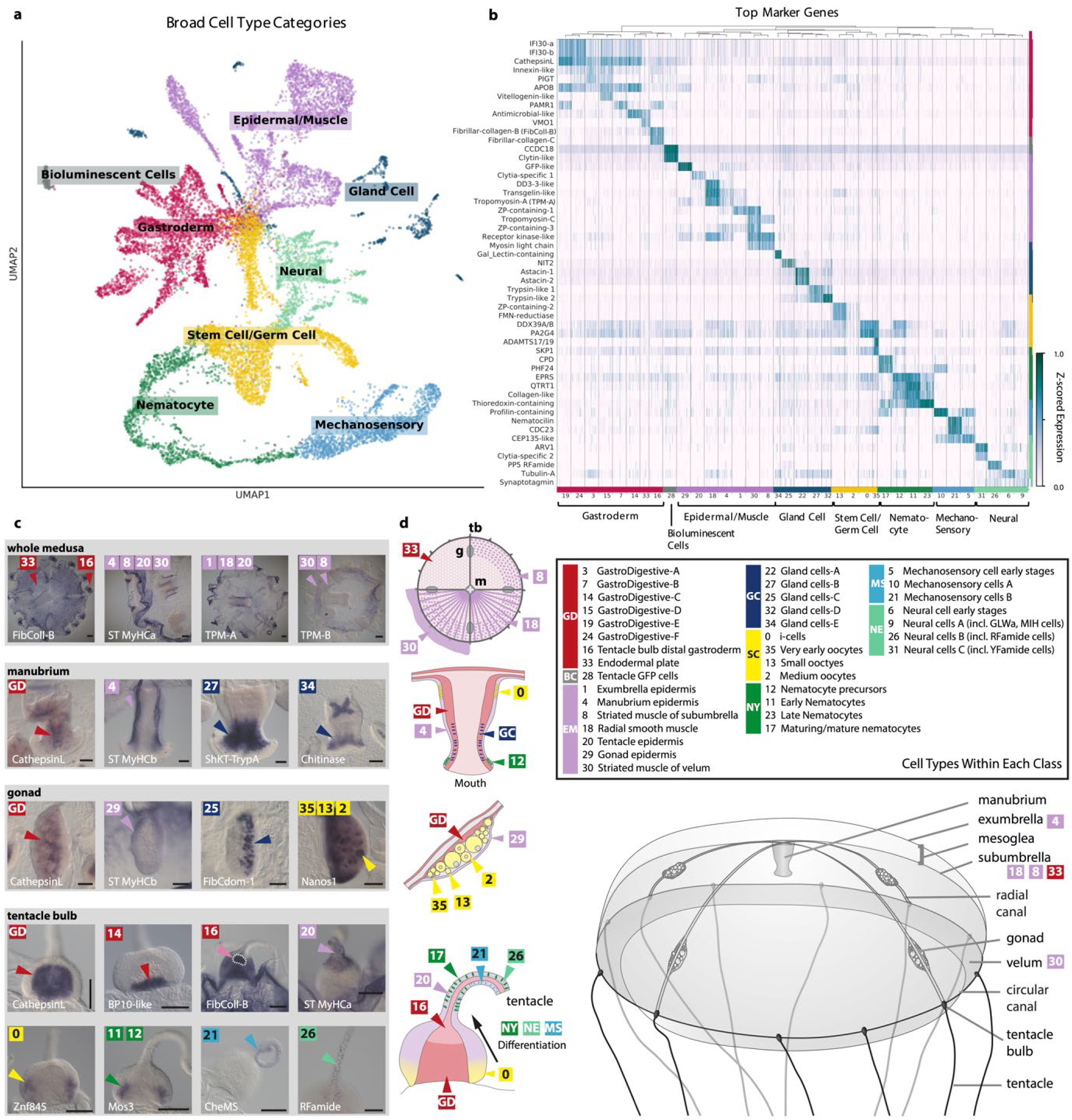
The *Clytia* Medusa Cell Atlas: a) 2D UMAP embedding of cells labeled by eight cell type classes. Class colors are retained in panels **b, c**, and **d. b)** Heatmap of top marker genes from the sequencing data with 36 Louvain clusters comprising the eight cell type classes. **c)** *In situ* hybridization patterns for a selection of cluster marker genes providing spatial location on the animal (comprehensive set in Supplementary Fig. 15). The label GD denotes general markers for gastroderm cell types. Scale bars: 100 µm. **d**) Schematics of *Clytia* medusa, manubrium, gonad, and tentacle bulb showing the main cell types. Abbreviations of cell class names -GD: gastroderm, BC: bioluminescent cells, EM: epidermal /muscle, GC: gland cells, SC: stem cell/germ cell, NY: nematocytes, MS: putative mechano-sensory cells, NE: neural cells. [[Code a,b](https://github.com/pachterlab/CWGFLHGCCHAP_2021/blob/master/notebooks/CellAtlasAnalysis/starvation_Analysis.ipynb)]

The 36 cell types (see Methods; Fig. 2b-d), were concordant between the two separate multiplexed experiments and robust to different transcriptome annotations (Supplementary Fig. 5,11,13). For some of them, cell type identity could be assigned on the basis of published information on gene expression in *Clytia* and/or of homologous genes in other animals, while for the others we performed *in situ* hybridization for selected marker genes (Fig. 2c; Supplementary Fig. 14,15; Supplementary Table 3). Previously known cell types apparent in our data included i-cells^33^ and nematocytes at successive stages of differentiation^34–36^, as well as oocytes^37^, gonad epidermis, manubrium epidermis, and bioluminescent cells in the tentacles that each express specific endogenous green fluorescent proteins (GFPs)^38^.

*In situ* hybridization for a selection of diagnostic muscle cell type genes allowed us to describe cell types making up the smooth and striated muscles, for instance, distinguishing the striated muscle cells lining the bell (subumbrella) and velum (Fig. 2c,d; Supplementary Fig. 15)^23,27^. Within known cell types, clustering revealed an unappreciated degree of cell heterogeneity, yielding novel subtypes. For example, six types of gastro-digestive (GD) cells, characterized by shared expression of enzymes associated with intracellular digestion, such as CathepsinL^39^, could be distinguished within the gastrodermis. These include GD-C (cluster 14) with striking expression patterns at the base of each of the gastrovascular digestive territories: stomach, gonad, and tentacle bulbs (Fig. 2c,d; Supplementary Fig. 14, 15). Two other gastrodermal clusters could be mapped spatially to distinct sites: the endodermal plate (cluster 33) or the tentacle-bulb junction (cluster 16). Markers for these two clusters, and for GD-C, included genes implicated in mesoglea (jelly) production, such as fibrillar collagens, indicating that these cell types contribute to shaping the medusa. Digestive gland cells fell into five types expressing different mixtures of enzymes for extracellular digestion, showing overlapping distributions spanning the mouth, stomach, and gonadal compartments of the gastrovascular cavity. Four broad clusters corresponding to neural cells each appeared to represent mixed populations, and could be subdivided by further analyses to define 14 likely subpopulations of neurons (see below). In addition, we uncovered cell types previously unknown in *Clytia*, such as putative mechanosensory (MS) cells expressing homologs of mechanosensory apparatus components characterized in vertebrate hair cells such as Harmonin and Whirlin^40–42^ (Fig. 2a-d, clusters 5, 10, 21; Supplementary Fig. 14,15). Expression of these genes was also associated with cluster 17 (assigned as maturing/mature nematocytes) consistent with assembly of the cnidocil mechanosensory apparatus of nematocytes^43^. In parallel, the highly characteristic marker genes of the stinging capsule (nematocyst), such as the minicollagens, downregulate^34,36^. Expression of marker genes for MS cells were detected in the tentacle bulb ectoderm and in cells positioned mainly on one side of each tentacle, opposite the row of oriented mature nematocytes (Fig. 2c,d; Supplementary Fig. 15).

A remarkable feature of the *Clytia* medusa is that it constantly generates many cell types, notably neural cells and nematocytes from prominent i-cell pools in the tentacle bulb epidermis^35^. The gland cell types may also be generated from i-cells, for instance from the pools at the base of the manubrium and in the gonad, as well as by self-renewal and position-dependent transdifferentiation as reported in *Hydra*^*8*^. Within our dataset, we expected to be able to capture dynamic information relating to the development of i-cell derived cell types, similar to that extracted from Hydra polyp single cell transcriptome data^8^. Indeed, our cell atlas embedding revealed neuronal and nematocyte populations connected to the i-cell population^3,8^ (Fig. 2a; Supplementary Fig. 14, 15), likely corresponding to differentiation trajectories^34,35^. To identify known and novel markers for nematocyte and neuronal development, we assigned pseudotime values to the cells and ranked genes in each trajectory (Fig. 3a, see Methods). This revealed trajectories consistent with both these populations deriving from i-cells^8^ (Fig. 3a). Examination of the nematocyte trajectory revealed the temporal expression of numerous genes previously not associated with this process, including developmental genes of interest, such as *Znf845* and *Mos3*^*44*^. *Genes previously known to be expressed during nematogenesis, such as Dkk3* and *Mcol3/4*, were also present at the expected times, providing an internal validation (Fig. 3b; corresponding expression domains in Supplementary Fig. 14, 15). Genes expressed during neuronal development included those encoding *bHLH, Sox*, and other transcription factors with potential roles in neurogenesis or fate specification, and numerous other genes of interest in neuronal development, such as cell adhesion molecules^35,45–47^ (Fig. 3c; Supplementary Table 5). Many other marker genes in both trajectories are uncharacterized and potentially *Clytia*-specific (Supplementary Table 5).

**Figure 3:**
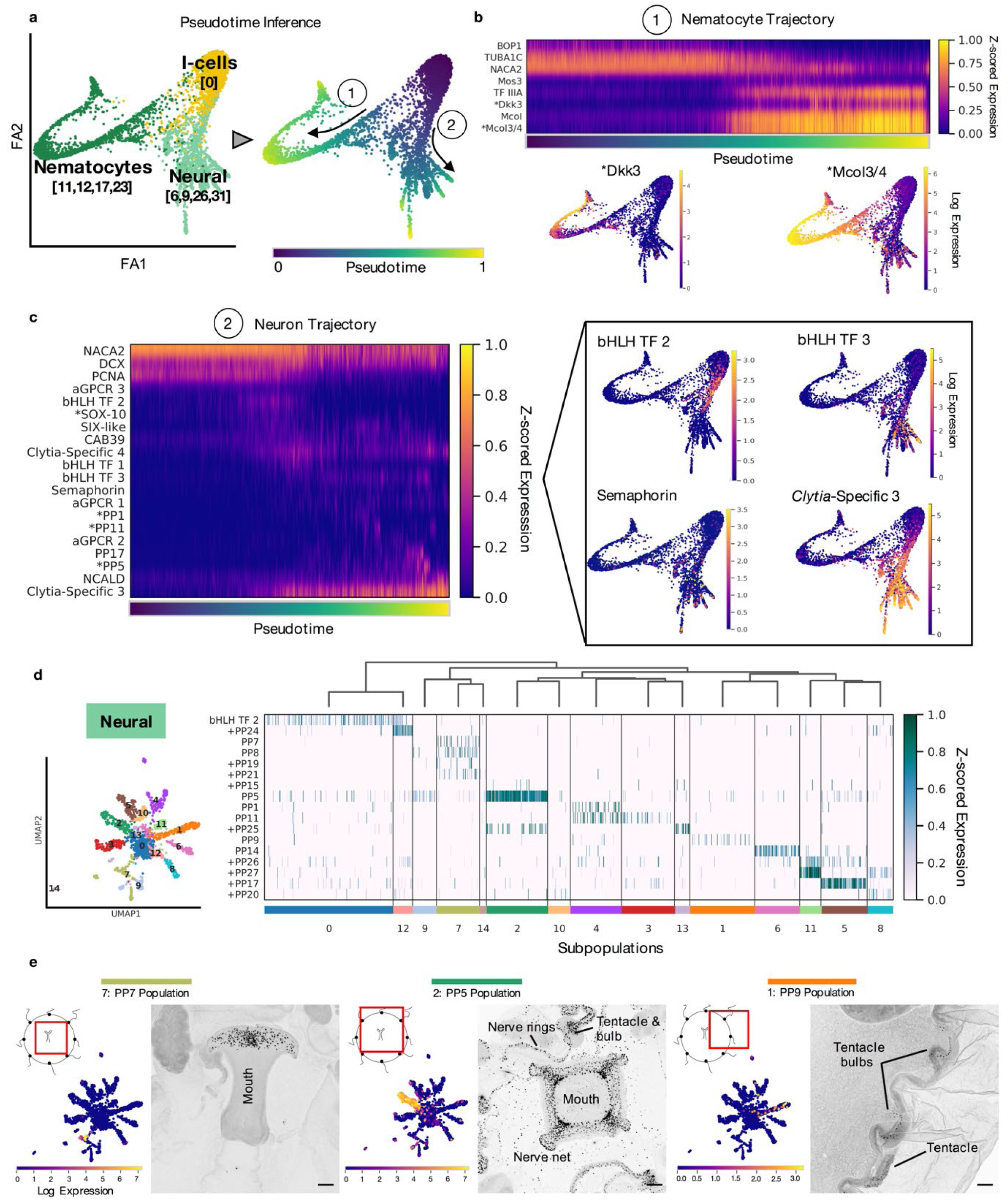
Developmental and Hierarchical Analysis of Neural Cell Types: a) Pseudotime trajectories for differentiating cells, **b)** Heatmap of expression of dynamic genes in nematocyte development over pseudotime. The embeddings below the heatmap show gene expression of known markers for nematocyte development along the trajectory. **c)** Heatmap of expression of dynamic genes in neuron development over pseudotime, selected from a ranking of genes according to neural trajectory (see Methods). ^*^ denotes previously reported genes. Embeddings on the right-hand side show expression of selected genes along the pseudotime trajectory. **d)** Subclustering of the ‘Neural’ group, with expression of predicted neuropeptides shown in the heatmap. + denotes previously unreported neuropeptides. **e)** Expression of neuropeptides, and corresponding *in situs* for the neural subpopulations they mark. Scale bar is 100µm. [[Code a-c](https://github.com/pachterlab/CWGFLHGCCHAP_2021/blob/master/notebooks/CellAtlasAnalysis/pseudotime_Analysis.ipynb)] [[Code d,e](https://github.com/pachterlab/CWGFLHGCCHAP_2021/blob/master/notebooks/CellAtlasAnalysis/neuronSubpop_Analysis.ipynb)]

Cnidarian nervous systems represent both valuable points of phylogenetic comparison and tractable platforms for systems neuroscience^3,8,20^. However, the molecular heterogeneity of neural cell types remains largely unexplored, particularly in the more complex medusa forms. Neuropeptides are thought to be the dominant neurotransmitters in cnidarians^48,49^ but are challenging to identify due to rapid sequence evolution^50,51^. In conjunction with sequence-based analysis, we were able to identify ten new likely neuropeptides on the basis of their inclusion as marker genes for the four basic neural clusters (6, 9, 26 and 31 in Fig. 2b,d), increasing the number of predicted *Clytia* neuropeptides to twenty-one (Supplementary Table 3). Our pseudotime ranking revealed that many of these predicted neuropeptides mark the later stages of neural cluster trajectories, likely defining distinct, mature neural subpopulations (Fig. 3c). We extracted and re-clustered the neural supergroup (‘Neural’, Fig. 2a,b) to characterize neural subtypes. This distinguished 14 subpopulations of neurons as well as a progenitor population, expressing cell cycle and conserved neurodevelopmental genes (sub-cluster 0, Fig. 3d). Strikingly, these neuronal subpopulations can be defined by combinatorial neuropeptide expression, often with a distinct and identifying neuropeptide (Fig. 3d). *In situ* hybridization for markers of neuronal subpopulations indicated that some have highly specific spatial locations while others are more broadly distributed across the animal (Fig. 3e). For example, the pp7+ subpopulation is located around the rim of the mouth and the pp9+ population is located in the tentacle bulbs and tentacles, while the pp5+ population is widely distributed across the tentacles, bulbs, nerve rings, subumbrella, and mouth (Fig. 3e). In comparison to neuropeptides, genes involved in classical chemical neurotransmission were not obvious markers for subpopulations in our dataset. There were, however, a number of interesting subtype-specific genes potentially involved in neurotransmission, for example, a possible nitric oxide synthase, and a member of the Slc6 family of transporters (Supplementary Table 5).

### Response to Starvation Across the Cell Atlas

To assess the transcriptional impact of starvation, we mapped individual cells to their corresponding control or starved labels, and placed the starvation-induced transcriptional changes within the context of the cell atlas, asking whether these changes resulted in new cell types, or shifts within a type, i.e. in cell state. As there are ∼5x fewer total cells in a starved animal (see Methods), we first asked whether there were significantly different numbers of cells per cluster between control and starved conditions. We found that only one cluster had a significant difference (cluster 11, early nematocytes; Supplementary Fig. 3), suggesting a nearly uniform reduction across cell types in the starved condition. In contrast, the distribution of cells from control versus starved animals across the atlas embedding showed dramatic shifts in the local density of cells from the two conditions within most clusters (Fig. 4a).

**Figure 4:**
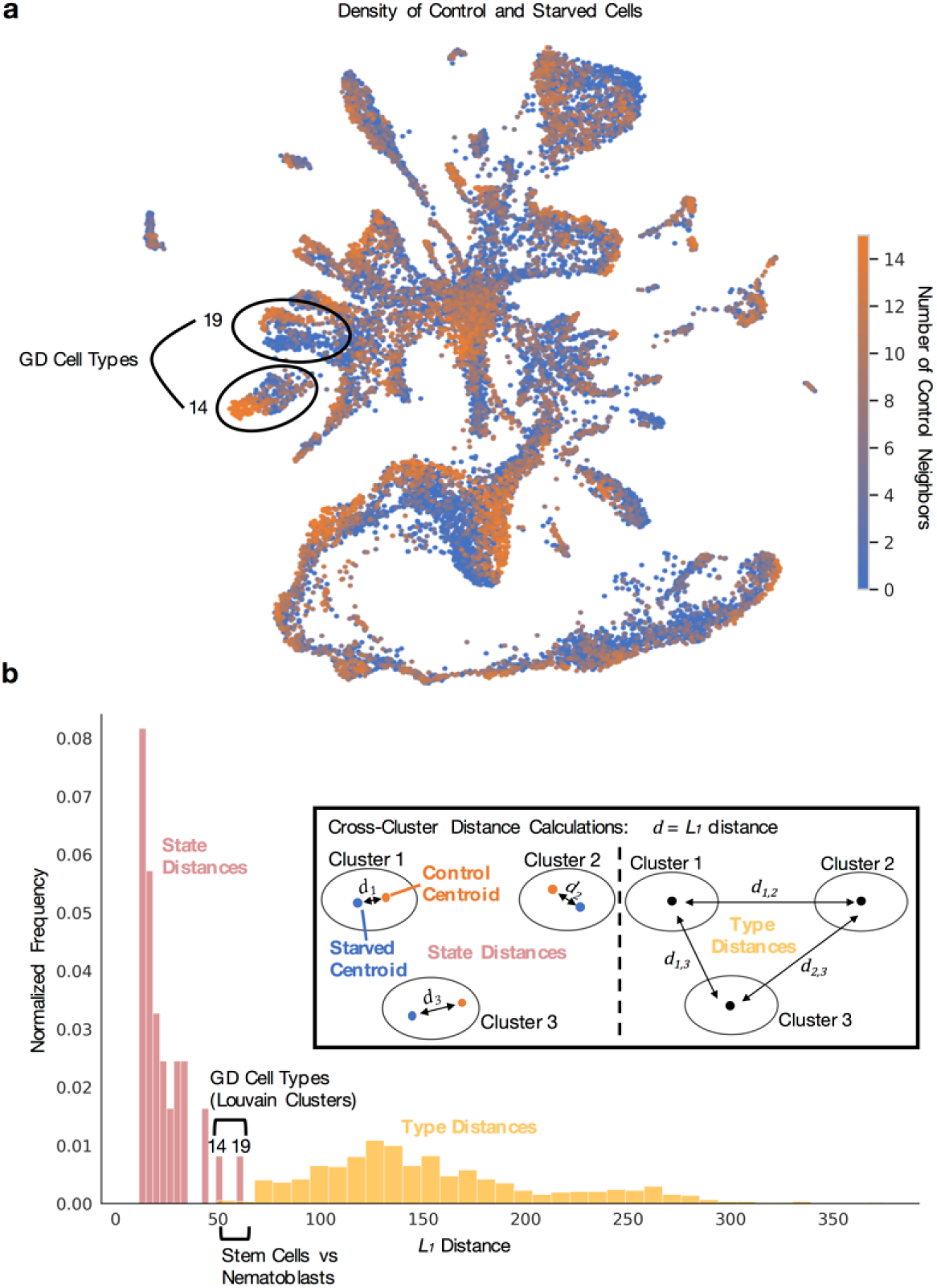
Starvation-Induced Shifts in Cell Types: a) Density map of fed and starved cells across cell types, with each cell colored by the number of its 15 nearest neighbors that are control cells. Clusters 14 and 19, with particularly dense and separated control/starved regions are circled. [[Code](https://github.com/pachterlab/CWGFLHGCCHAP_2021/blob/master/notebooks/CellAtlasAnalysis/starvation_Analysis.ipynb)] **b)** Histogram of *L*_*1*_ distances between centroids of control and starved cells within cell types, versus pairwise *L*_*1*_ distances between centroids of the 36 cell types. Clusters 14 and 19, with largest internal distances are highlighted, and clusters with small pairwise distances also noted for comparison. Inset illustrates inter- and intra-*L*_*1*_ distance calculations. [[Code](https://github.com/pachterlab/CWGFLHGCCHAP_2021/blob/master/notebooks/ComparativeDistanceAnalysis/allDistanceCalculations.ipynb)]

To determine whether starvation had generated new cell types (as defined operationally by the clustering), we asked how the distances between control and starved cells within clusters compared to the distances between clusters. As a metric, we used the *L*_*1*_ distance (see Methods), i.e. the sum of the absolute differences between centroid coordinates in PCA-reduced space. We found that the *L*_*1*_ distances between control and starved cells within a cell type, versus between cell types, formed nearly non-overlapping distributions (Fig. 4b, see Methods). This suggests that, overall, *Clytia* do not generate new cell types in response to starvation, but rather respond with shifts in cell state, and that their cell type repertoire is well represented by the original clusters. However, the impact of starvation was variable across cell types, as reflected by the range of internal (state) distances (Fig. 4b). Starvation produced the largest perturbations in cells of the gastrovascular system, causing control-vs-starved distances large enough to overlap with the smallest inter-type distance, i.e. that between the stem cells and nematocyte precursors (Fig. 4b). This distinction between state shifts and type was also clearly visible in the lack of overlap between the distributions of inter- and intra-cluster distances within the second multiplexed experiment (Supplementary Fig. 5e). Though classification and distinction of cell state and type is a complex task^52^, this analysis, based on relative distance in transcriptional space, provides a quantitative basis for delineation of type/state effects that may be useful in other contexts.

To characterize gene-level responses underlying these starvation-induced shifts, we then asked if responses are shared or unique across the cell types and compared the extent of the responses, in terms of gene quantity and expression level, across the atlas. For each cell type, we collected genes that were differentially expressed under starvation (“perturbed genes”; see Methods, Fig. 5a). For a high-level view of the general functions and processes affected by starvation, and their cell type specificity, we clustered perturbed genes into apparent “gene modules”^53^ by their pattern of co-expression across cells (Fig. 5a; see Methods). We assigned putative functions to these gene modules through GO term enrichment, giving a global view of affected processes (Supplementary Fig. 16), and examined the distribution of cell types across modules by asking in how many cell types is a given gene a perturbed gene (Fig. 5b). We found that certain gene modules were broadly shared across cell types, while others were almost entirely cell type-specific (Supplementary Fig. 16). Striking examples include gene module 5, which is enriched in proteolytic genes (Fig. 5c, Supplementary Fig. 16) and has shared expression across multiple GD cell types (Fig. 5c). Notably, there is also divergent gene expression between GD types (Fig. 5c). In comparison, gene module 3 is largely composed of early oocyte gene expression (∼70%), and is enriched in cell cycle and developmental genes (Fig. 5d). These modules thus provide an overview of which processes affected by starvation are shared across cell types and reveal divergent expression potentially reflecting different motility^19^, locations, or as-yet-undescribed functional differences between the cell types (Supplementary Fig. 17).

**Figure 5:**
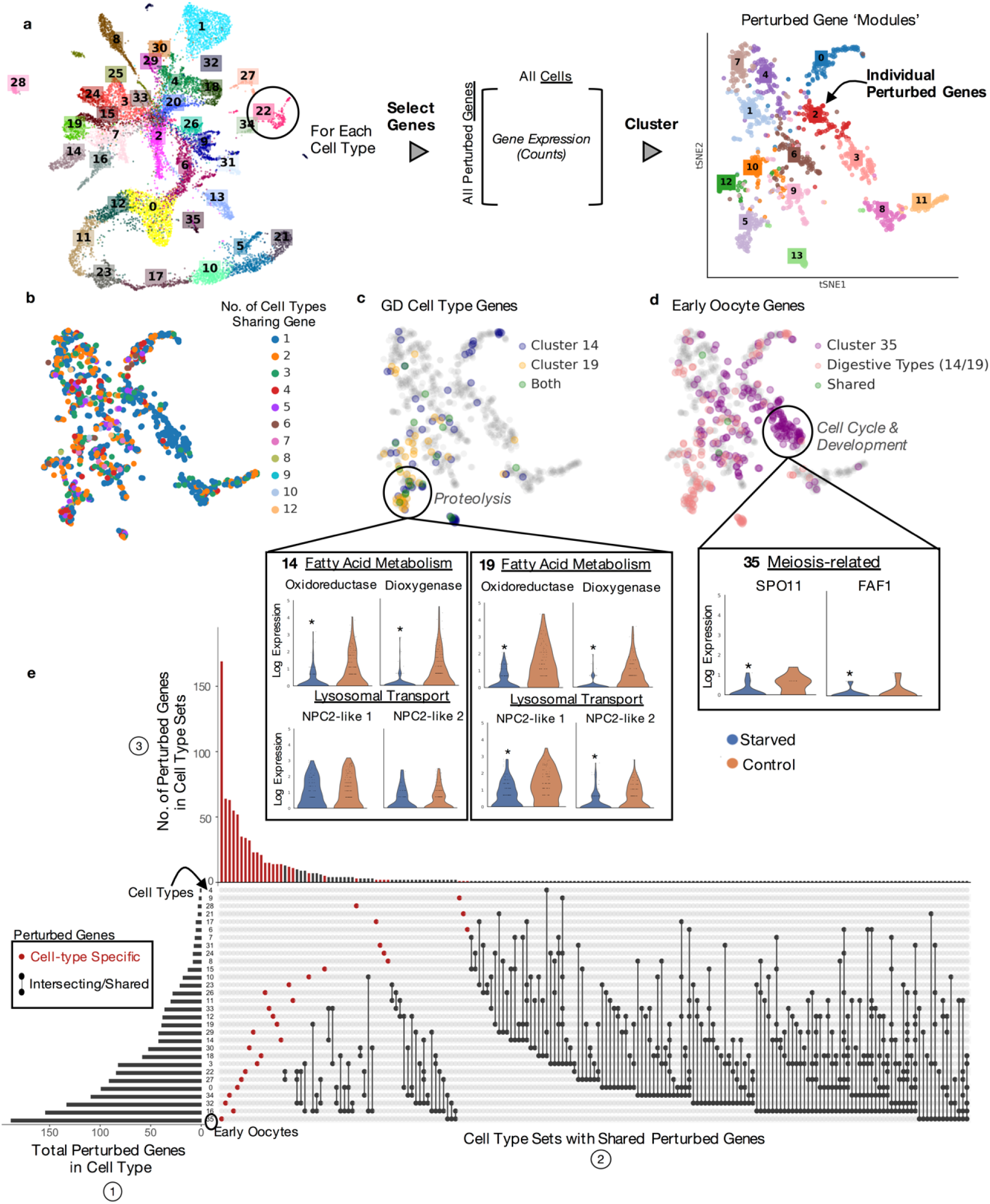
Perturbation Responses in the Oocyte and Digestive Cell Types: a) Workflow for extracting ‘perturbed’ genes per cell type and clustering on genes to extract co-expressed groups or ‘modules’ of genes (Supplementary Fig. 16). **b)** Each gene in module embedding colored by number of cell types in which the gene was differentially expressed under starvation, **c)** Visualization of locations of perturbed genes (among the embedded modules) of the two highly perturbed GD cell types (clusters 14, 19) and **d)** early oocytes (cluster 35). Violin plots showing expression profiles for several perturbed (differentially expressed) genes in ‘functional’ categories of interest. ^*^ *p*-value < 0.05. Horizontal lines show quartiles and width of violins denote density of points in that region. **e)** UpSet plot visualization for intersecting sets of ‘perturbed’ genes. The barplot in 1) shows the number of genes that are differentially expressed under starvation (‘perturbed’ genes) for each cell type. This is the cardinality or size of each cell type’s set of perturbed genes. 2) The connected dots represent intersections (overlaps) between the sets of each cell type’s perturbed genes. Genes with starved differential expression in only one cell type are denoted by the red, unconnected dots. 3) The top bar plot is the number of perturbed genes within these sets of shared perturbed genes, i.e. the cardinality of the intersections in 2. [[Code a-d](https://github.com/pachterlab/CWGFLHGCCHAP_2021/blob/master/notebooks/CellAtlasAnalysis/starvation_Analysis.ipynb)] [[Code e](https://github.com/pachterlab/CWGFLHGCCHAP_2021/blob/master/notebooks/CellAtlasAnalysis/deSeq2Analysis_StarvationResponse.ipynb)]

To examine how individual perturbed genes are distributed across cell types, we visualized, for each cell type, how many perturbed genes it had, and how many of these genes are unique versus shared with other cell types (Fig. 5e). We found a large number of perturbed genes (∼72%) were cell type specific (Fig. 5d). For the most perturbed cell types, we examined whether the state shifts that we had observed were due to changes in a large or small number of genes, and how highly these genes were expressed. Consistent with the marked shrinkage of the gonads during starvation treatment (Fig. 1), early oocytes contained the highest number of perturbed genes, which were spread across many gene modules (Fig. 5d). In contrast, the GD cell types had fewer perturbed genes that were expressed at higher levels and localized to more specific gene modules (Fig. 5c), highlighting the diverse logic employed by cell types under starvation (Supplementary Fig. 18).

In accordance with these distinct responses in GD cells and oocytes, comparison of the cellular organization of gonads from control and starved medusae revealed major reorganization of both the gastrodermis and the oocyte populations (Fig. 6). Most strikingly, the population of mid-sized, growing oocytes, which progress daily through vitellogenesis in conditions of normal feeding^29^, was largely depleted following starvation, leaving a majority of pre-vitellogenic oocytes (Fig. 6a). A sparse population of large oocytes in starved gonads likely results from growth of a minor subpopulation of oocytes fueled by recycling of somatic tissue and oocytes (disintegration of smaller oocytes visible in Fig. 6a, asterisks). Consistently, GD cells in many parts of the gonad lost their regular epithelial organization and, despite the absence of any external food supply, showed evidence of active phagocytosis involving variably-sized vesicles (arrows in Fig. 6a). Changes in organization and activity of the gonad gastrodermis were also evident from comparison of expression patterns for the GD cell-expressed genes CathepsinL and Nucleoside hydrolase, and a ShKT and trypsin domain protease (*ShKT-TrypA*) expressed in gland cell types A and B, which is down-regulated during the starvation treatment (Fig. 6b). Shifts between gonad gastrodermis organization and transcriptional profiles induced by starvation thus accompany activation of tissue autodigestion programs, and also the mobilization of GD cells (termed MGD for Mobilizing Gastro Digestive cells^19^) from the gonad through the gastrovascular canal system, which has been observed both under conditions of starvation and during regeneration of the feeding organ^19^.

**Figure 6:**
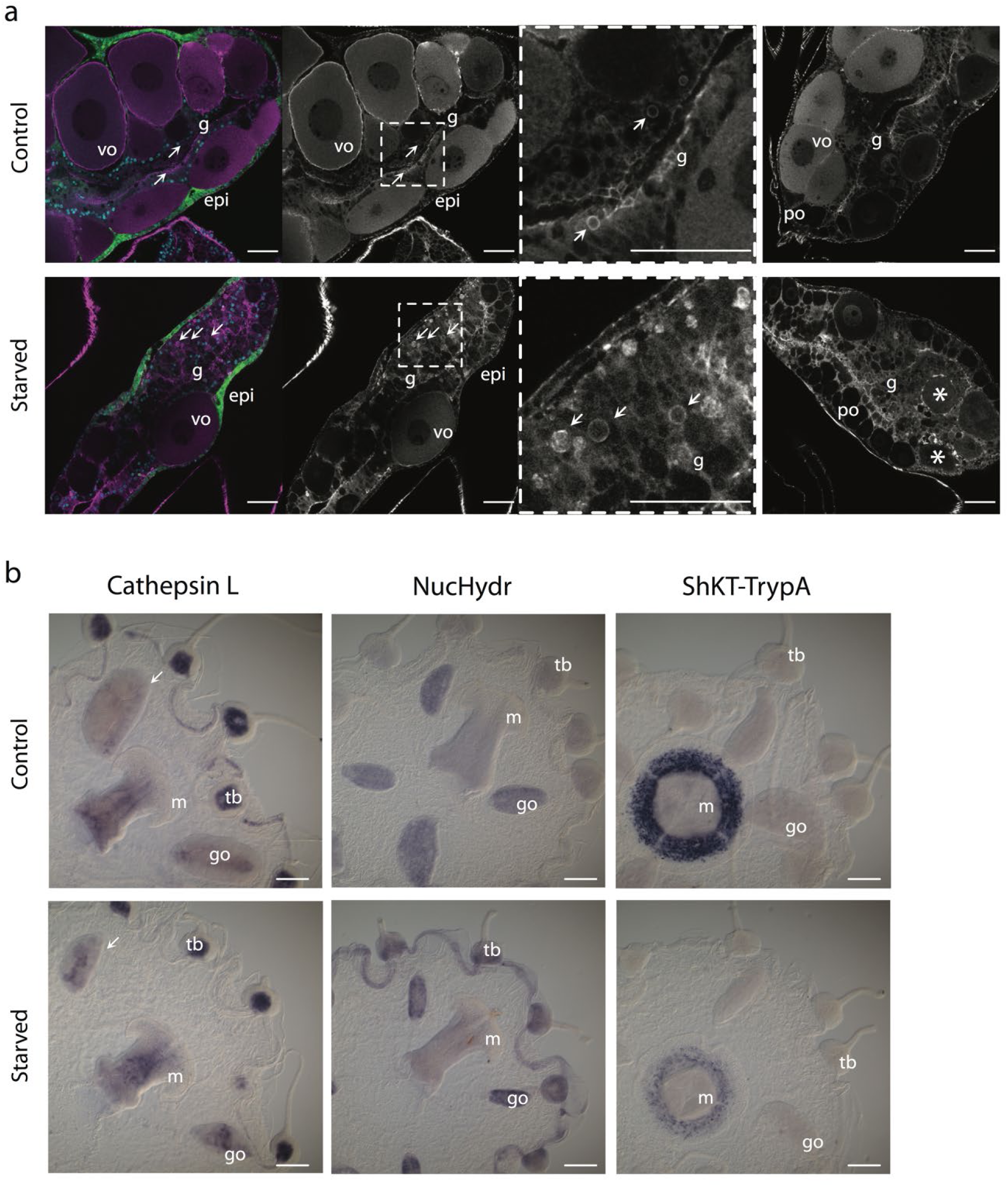
Perturbations to gastroderm cell types in response to starvation. a) Confocal sections through gonads from control and starved medusae with cell morphology revealed by phalloidin staining of cell boundaries (magenta/grey). The first panel of each row shows co-staining of nuclei with Hoechst (blue) and endogenous GFP4 (green) in the outer epidermis (epi). Vitellogenic oocytes (vo) are largely absent following during starvation, leaving a majority of pre-vitellogenic oocytes (po). The gastroderm (g) is heavily reorganized, with evidence of active phagocytosis (vesicles arrowed) and disintegrating oocytes (asterisks). The third panel in each row is a higher magnification of the boxed area in the second panel, and the fourth panel shows a second example gonad for each condition. Scale bars represent 50µm. **b)** *In situ* hybridization of cell-type specific gene expression affected by starvation. The general GD cell marker CathepsinL confirms extensive reorganization of the gastroderm in starved animals, especially in the oocyte-depleted gonad (arrows) ; Nucleoside hydrolase (NucHydr), a GD-B marker is detected more strongly and is more widespread following starvation, while the gland cell A/B gene ShKT and trypsin domain protein A (ShKT-TrypA) is down-regulated. Gonads (go), manubrium (m), tentacle bulbs (tb). Scale bars represent 200µm.

## Discussion

The *Clytia* medusa single-cell atlas presented here is an important addition to the growing number of single-cell atlases across the animal tree of life. It provides the first cell-level transcriptomic characterization of a pelagic medusa stage, the most complex of the life cycle forms within the large and diverse phylum Cnidaria. Reflecting this complexity, we found greater cell type diversity in the *Clytia* medusa than in its polyp-only hydrozoan cousin *Hydra*^*8*^. *The outer, epidermal body layer could be sub-divided into seven clusters encompassing all of the described Clytia* muscle types, including two types of fast-contracting striated swimming muscle^23,27^. Rich diversity was also uncovered in the inner gastroderm layer, which is elaborated in the medusa into distinct digestive compartments (mouth, stomach, gonad, and tentacle bulb) and also generates the thick mesoglea (jelly) characteristic of the medusa form. Of the eight gastroderm cell clusters, four could be mapped to distinct sites by marker gene *in situ* hybridization including three likely involved in mesoglea/ECM production and thus in modulating the medusa structure. A total of six gastrodermal clusters represent cell types with a major role in intracellular digestion based on their shared set of marker genes (Fig. 2b; Supplementary Table 5). These include four that are widely distributed across the digestive compartments, suggesting overlapping functional specialization. Our starvation experiment analyses revealed that these clusters were maintained operationally as distinct ‘cell types’ rather than ‘cell states’ between the two extreme conditions tested, but we cannot rule out that responses to other environmental or physiological perturbations may reveal plasticity between these clusters.

In addition to the epithelial cell types of the epidermis and gastroderm, our single-cell atlas supports the presence of an interstitial stem cell (i-cell) population in *Clytia* providing a similar set of somatic cell types to that described in *Hydra*, as well as the germ cells^8,33^. Our pseudotime analyses provided transcriptional signatures of the progressive stages of nematogenesis and neurogenesis from i-cells that will guide future studies of their developmental regulation. The newly-described putative mechanosensory cells^40,41^ likely also derive from i-cells. We uncovered 14 mature neuronal subtypes in *Clytia*, which is similar to the number reported in *Hydra* and *Nematostella*^*3,8*^. However, it is likely that further heterogeneity exists within these 14 subpopulations. These subpopulations could be defined by combinatorial neuropeptide expression, with some located in specific anatomical regions, and others distributed across the medusa (Fig. 3d-e). How molecular cell type maps to function both within and across body parts, the roles of these defining peptides as primary transmitters and/or neuromodulators, and the uses, if any, of classical, small-molecule neurotransmission, remain unknown. Moving forward, with this cell atlas as the foundation, the ability to perform whole-organism, multiplexed scRNAseq, in combination with emerging genetic tools and advantageous life history traits, makes *Clytia* a powerful, tractable platform for high-resolution systems biology.

This work additionally serves as a case study in using multiplexed single-cell transcriptomics to assess cellular responses to whole organism perturbations, and provides a guide for deployment in other organisms. We anticipate WHAM-seq will be impactful for researchers studying various biology, from developing embryos to organoids to non-model organisms. The techniques for multiplexed experimentation that underlie this study are also well suited to large-scale perturbation studies in other marine organisms given the sea-water compatible workflow. As sequencing costs drop and cell throughput in scRNA-seq grows, WHAM-seq should also become tractable for larger, more complex systems. Moreover, the lack of library-induced batch effects demonstrates how large-scale experiments can be conducted without introduction (or minimizing introduction) of confounding factors from multiple experiments, which can be highly non-linear and difficult to account for^13^. The second perturbation dataset also demonstrates how batch effects are reduced within multiplexed experiments (see Methods; Supplementary Fig. 10).

The fully reproducible and usable framework we have presented, which includes code that can be run on a laptop or for free in the cloud (see Code Availability), will further assist in extending this expression-based analysis to other organisms. By relying on expression, our strategy reduces the reliance on prior gene functional annotation, using specificity of expression (gene ‘activity’) to identify genes of interest, allowing for targeted annotation. ‘Activity’ includes determination of strong diagnostic markers for cell type definition, cell type specific and shared transcriptional responses to starvation, and ‘modules’ of co-expressed genes underlying these responses. The extent of these expression-based changes additionally highlights areas of the organism’s biology that are strongly or uniquely affected by a perturbation, in this case, honing in on the large scale downregulation of gene expression in two digestive gastroderm cell types and severe disruption of oocyte development under starvation. Together, this “reverse genomics” approach dramatically lowers the barriers for working with non-traditional models, and affords opportunities to match uniquely-suited organisms to specific questions. Moving forward, the combination of scRNA-seq and other sequencing-based genomics techniques with multiplexing and annotation-agnostic analyses, could foster comprehensive high-resolution molecular studies of diverse organisms and their responses to numerous environmental perturbations.

## Methods

### Animal culture and experimental setup

#### Starvation

Culture of the *Clytia* life cycle was carried out as previously described^54^, with some modifications to the tank design. The system used in this study to culture polyps uses zebrafish tanks (Pentair), with polyp slides held in glass slide racks (Fisher, cat#02-912-615). Medusae used in this experiment were raised in 4L beakers with a circular current generated by stirring with a constant speed 5rpm DC motor (Uxcell), attached to the lid of a multiwell tissue culture plate. Artificial sea water for culture and experiments was made using Red Sea Salts (Bulk Reef Supply, cat# 207077) diluted into building deionized water to 36ppt. Experiments used ∼1cm medusa of the Z4B strain.

For the experiment, baby medusae were collected overnight and then cultured together until they reached ∼1cm (about 2 weeks). Animals were fed once per day using 2-4 day old brine shrimp. Prior to the experiment, animals were split into two beakers. Feeding continued as before for one beaker, while the other was starved for 4 days. The “control” group was not fed on the day of the experiment. The 4 day time point was chosen as animals show strong phenotypic changes (Fig. 1) but it is far from their survival limit following starvation, as *Clytia* medusae will survive for more than 3 weeks with no food.

#### Stimulation

Rearing of the medusae prior to the experiment was performed as described above. On the day before the experiment, each *Clytia* medusa (3-5 weeks old) was placed in a separate container (∼150 mL seawater), which was covered with foil and moved to the experimentation area to acclimate overnight in the dark. The morning following overnight acclimation, lights were turned on for ∼2.5 hours to allow for spawning to complete^55^. Each animal was then given repeated bouts of stimulation over a period of 30 mins, with each stimulus administered every 2 minutes. 100 uL of each stimulant (150 mM KCl, DI water, or seawater (SW) as a control) was gently added just below (or just above for KCl) each medusa by pipette. (Supplementary Fig. 4). Stimuli were chosen based on their ability to reliably induce crumpling behavior, a protective response in which the bell is drawn in towards the mouth using the radial muscle^56^.

### Single-cell suspension and multiplexing

For the starvation experiment, animals were washed with hypertonic PBS (500 mM NaCl, 2.7 mM KCl, 8 mM Na_2_HPO_4_, and 2 mM KH_2_PO_4_, pH 7.4) by serial transfer from seawater through three successive containers each with 150 mL hypertonic PBS, to prepare cells for fixation and to avoid precipitation of seawater salts in methanol. The animals were then thoroughly homogenized with a douncer. After homogenization, cells were collected by centrifugation at 500 x *g* for 5 minutes and resuspended in 100 uL hypertonic PBS. Cells were then fixed by addition of 400 uL ice-cold methanol and stored at −80C until sample indexing and library preparation. Each sample was labeled according to the ClickTag labeling procedure described previously^14^. ClickTags used for each animal are outlined in Supplementary Table 3. Each sample was labeled with two distinct and unique ClickTags, and samples were then pooled after addition of the ‘blocking’ oligo. Fed and starved samples were counted on a Countess, both to estimate cell numbers per animal and to determine concentrations for 10X loadings. 200,000 cells/mL were counted for the starved sample and 1 M cells/mL for the control. All the starved cells and (an equivalent) 200,000 cells of the control were then pooled and loaded into two lanes of the 10X Chromium Controller with v2 chemistry. Sample tag libraries were separated and processed after an SPRI size-selection step as previously described^14^. cDNA samples were run on 2 lanes of HiSeq 4000 (two HiSeq 3000/4000 SBS 300 cycle kits) and tag libraries were run on 2 lanes of MiSeq (using MiSeq v3 150 cycle kits).

The same protocol was followed for fixing and labeling cells from the stimulation experiment. Each animal in this case was assigned one unique ClickTag and one ClickTag per condition (Supplementary Table 3). After the separation of the cDNA and ClickTag samples, ClickTags were added at a 3% final concentration to the cDNA samples sequenced on the HiSeq, in addition to separate sequencing on the MiSeq. The full protocol is described in the Supplementary Methods.

### In Situ Hybridization

Colorimetric *in situ* hybridization (ISH, Fig.2, 6b) was performed as previously^57^ with minor modifications. In brief, 2-week-old medusae (Z4B strain) were relaxed in 0.4 mM menthol in sea water and tentacles were trimmed prior to fixation in a pre-chilled solution of 3.7% Formaldehyde, 0.2% Glutaraldehyde in PBS on ice for 40 min. Specimens were then washed thoroughly with PBST (PBS + 0.1% Tween20), dehydrated in methanol stepwise and stored in 100% methanol at −20°C. Hybridisation (at 62°C for 72h) and washing steps were performed in a robot (Intavis AG, Bioanalytical Instruments) using 20X SSC pH adjusted to 4.7 throughout. Acetylation steps using 0.1M triethanolamine in PBST (2×5min) then 0.25% acetic anhydride in 0.1M triethanolamine (2×5min) followed by PBST washes (3×10min) were included before pre-hybridization to reduce probe non-specific binding. Incubation with Anti-DIG AP, 1:2000 in 1X blocking solution was performed for 3h before washing and the NCB-BCIP colour reaction at pH 9.5. Following post-fixation, washing and equilibration of samples in 50% glycerol/PBS, images were acquired using a Zeiss AxioImager A2. Fluorescent ISH in Figure 3 was performed as previously described^57^, with minor modifications. Briefly, Anti-DIG-POD (Sigma 11207733910) was used during the antibody step. Following overnight incubation in antibody, animals were washed with maleic acid wash buffer (MABT; wash buffer from Sigma 11585762001) 6 x 15 min and with TNT (Tris-HCl pH7.5, 0.15 M NaCl and 0.1% tween 20) 3 x 15 min. They were then incubated in TSA solution (PerkinElmer NEL749A001KT; ratio: 1.5 ul TSA, 250 ul diluent, 1.25 ul 10% Tween 20) for 20 min at RT, washed 3 x 15 min with MABT, and incubated with 1:100 dilution of streptavidin-594 (1 ug/ml; ThermoFisher, S32356) solution, before a final set of 3 x 20min washes in MABT. Animals were then imaged using an Olympus FV3000 confocal microscope.

Probes were generated by PCR from cDNA clones corresponding to our EST collection^58^ or from medusa cDNA; the Elav probe was synthesized as a gBlock by Integrated DNA Technologies (details in Supplementary Table 3). For probes against the pp9 marker gene and Elav, the T3 polymerase recognition site (AATTAACCCTCACTAAAGGG) was added to the 3’-end of the PCR product, or gBlock, respectively. Products were TOPO cloned (ThermoFisher cat#K280020) and sequence verified. All probes were labelled with DIG RNA labelling mix (Sigma-Aldrich 11277073910), and purified with ProbeQuant G-50 Micro Columns (GE HealthCare Life Sciences Cat# 28-9034-08).

### Confocal Microscopy

Visualization of cell morphology within the gonads of control and starved young adult female medusae (Z4B strain) by confocal microscopy was performed as previously^55^. Fixation used 4% EM-grade paraformaldehyde in 0.1M Hepes pH 6.9/50mM EGTA/10mM MgSO_4_/80mM Maltose/0.2% Triton X-100 for 2h at room temperature. Specimens were washed 3×15min with PBS/ 0.02% Triton and 3×5 min in PBS. Cell boundaries and nuclei staining was performed by overnight incubation in 1/50 rhodamine-phalloidin (Molecular Probes 1mg/ml) and 1/5000 Hoechst 33258 (Img/ml stock, Sigma Aldrich) in PBS. Samples were washed 3×15min with PBS /0.02% Triton and 3×5min with PBS and equilibrated in 50% PBS/ Citifluor (Citifluor AF1) before imaging using a Leica SP5 confocal microscope. Control medusae were fixed 24h after the last feeding while starved ones were fixed 4 days after the last feeding.

### Generation of Reference Transcriptome

Assignments to PANTHER database entries (version 11) were made using the ‘<u>pantherScore2.0.pl</u>’ script available from the database website (www.pantherdb.org) (ref: PMID: 30804569). Human -*Clytia* orthologs were assigned using the OMA program (ref: PMID: 31235654) as described in PMID: 30858591, and taken from the pairwise human / *Clytia* orthologs output, rather than the orthologous groups.

To ensure highly sensitive transcriptome alignment, a new transcriptome assembly for *Clytia* Hemisphaerica was generated from bulk RNA-seq data produced from *Clytia* medusae (organisms at the same life stage as in the single-cell experiments) ([link](https://www.ncbi.nlm.nih.gov/sra/ERX2868482%5Baccn%5D)). We used the Trinity^59^ *de novo* assembler, with default parameters, to generate a transcriptome ([link](https://data.caltech.edu/records/1824)) and the Cufflinks Cuffcompare utility^52^ to merge the Trinity assembled transcripts with any XLOC annotations from the MARIMBA v.1 (created on 05/30/2016, [link](https://data.caltech.edu/records/1830)) transcriptome assembly^21^. Then with the CD-HIT ([link](http://weizhongli-lab.org/lab-wiki/doku.php?id=cd-hit-user-guide)) clustering method, assembled sequences with at least 95% were clustered and only one representative sequence was kept for each cluster. With the published MARIMBA v.1 genome sequence as a reference (created on 5/30/2016, [link](https://data.caltech.edu/records/1828))^21^, the GMAP ([link](https://academic.oup.com/bioinformatics/article/21/9/1859/409207)) aligner converted the collapsed Trinity fasta records to gff3 coordinate file ([link](https://data.caltech.edu/records/1824)). Most differentially expressed genes found in single-cell RNA-seq data from this study were previously identified and annotated (Supplementary Fig. 13). This annotation was used for the pre-processing and quantification of single-cell RNA-seq described below. Protein sequences were then obtained by running Transdecoder^60^ with default settings for the Trinity transcriptome ([link](https://data.caltech.edu/records/1827)).

### Pre-Processing and Clustering of Sequencing Data

#### Initial Cell Ranger Demultiplexing for ClickTags

Initial demultiplexing of ClickTag libraries was done using output from with the 10X Cell Ranger pipeline, using Cell Ranger 3.0 *count* and *aggr* functions with the MiSeq ClickTag fastqs as input, and combining the counts from the two lanes using the denoted sample IDs. A ClickTag count matrix (ClickTag x cell) was generated by counting ClickTag barcodes that had high sequence similarity to the designed sequences using the Python fuzzywuzzy package to identify targets within Levenshtein distance 1. [[Code](https://github.com/pachterlab/CWGFLHGCCHAP_2021/blob/master/notebooks/Preprocessing/ClickTagDemultiplexing/cellRangerClickTagCounts.ipynb)]

We additionally quantified gene expression with the kallisto-bustools workflow^61^ to create a more integrated workflow, which reproduced concordant results, described below.

#### Initial Cell Ranger Demultiplexing and Clustering for cDNA

Initial processing of starvation cDNA libraries was performed with the 10X Cell Ranger pipeline, using Cell Ranger 3.0 *count* and *aggr* functions to align and quantify the HiSeq reads, and combining the counts from the two lanes using the denoted sample IDs. This was followed by filtering cells for the high-quality cells chosen during ClickTag analysis, in addition to filtering cells by thresholding the rank-UMI vs cell barcodes plot. Values were log1p normalized, mean-centered, and scaled for downstream dimensionality reduction and visualization using Scanpy^62^. [[Code](https://github.com/pachterlab/CWGFLHGCCHAP_2021/blob/master/notebooks/Preprocessing/cDNAFiltering/filterStarvCells_ClickTags.ipynb)]

We then conducted Louvain clustering^63^ on the data mapped to a lower dimensional space, by applying PCA to the expression data filtered for highly variable genes, initially using Scanpy’s filter_genes_dispersion on only the log-normalized data. This resulted in the identification of 36 clusters (Fig. 2b; Supplementary Fig. 11), which we also refer to as cell types. The marker genes were selected by analyzing the top 100 markers extracted by Scanpy’s rank_genes_groups using default settings (*p*-values adjusted with the Benjamini-Hochberg method for multiple testing). The clusters were annotated and validated with marker genes previously identified in the literature, and manually categorized into the eight classes in Fig. 2a based on the marker gene patterns and their functional annotations. [[Code](https://github.com/pachterlab/CWGFLHGCCHAP_2021/blob/master/notebooks/CellAtlasAnalysis/cellRangerClustering_Starvation.ipynb)]

#### kallisto bustools for Demultiplexing and Clustering: Standardization of Workflow

In order to integrate and update the analysis using a platform with streamlined ClickTag demultiplexing and count matrix generation workflows, we used the kITE demultiplexing protocol ([link](https://github.com/pachterlab/kite)), which is based on the kallisto-bustools workflow and is described in the ClickTag demultiplexing protocol^14^. Briefly, MiSeq reads are aligned to possible tag sequences (Hamming distance 1 away from designed oligo sequence whitelist) by building a kallisto index and pseudoaligning reads to this index. Counts for these sequences were then collapsed into counts for their respective ClickTags, creating a cell-by-tag count matrix. We used Louvain clustering of cell barcodes based on the observed ClickTags, to filter for clearly delineated cells (clusters strongly marked by the individual’s two corresponding tags) and exclude sample doublets^14^. We also followed similar pre-processing to standard cell-by-gene workflows, using the inflection point in rank-UMI vs cell barcodes knee plots to filter cell barcodes based on their tag UMI counts. [[Code](https://github.com/pachterlab/CWGFLHGCCHAP_2021/blob/master/notebooks/Preprocessing/ClickTagDemultiplexing/kallistobusFedStarvedClickTagsProcessing.ipynb)]

For the stimulation experiment we concatenated sequencing data from the MiSeq and HiSeq as input to the previously described kallisto-kITE workflow. With the same clustering procedure we selected cell barcodes in clusters with strong overlapping expression of both individual and condition ClickTags. [[Code](https://github.com/pachterlab/CWGFLHGCCHAP_2021/blob/master/notebooks/Preprocessing/ClickTagDemultiplexing/stimClickTags.ipynb)]

To standardize the cDNA analysis workflow, we re-processed the starvation data using the kallisto-bustools workflow to generate gene count matrices for each lane, which were then concatenated. Cells were also filtered based on the ClickTag analysis. Values were log1p normalized, mean-centered and scaled, and filtered for highly variable genes using the same procedure described above for downstream dimensionality reduction and visualization (e.g. PCA) using Scanpy. We found that with the kallisto-processed data (with the same Cell Ranger clustering applied to the cells), the top 100 markers for each of the 36 clusters determined with Scanpy’s rank_genes_groups function (Supplementary Table 5) (using the nonparametric Wilcoxon test) overlap with markers in the Cell Ranger expression data, verifying that the cluster labels were concordant (Supplementary Fig. 12). We used the PAGA graph abstraction method^30^ to generate an underlying graph representation of the connectivity between cells (determining connectivity by the number of inter-edges between cell groups compared to the number of inter-edges under random assignment). We then generated a 2D UMAP^31^ embedding initialized with the PAGA graph structure for cell atlas visualization (Fig. 2a) using Scanpy. For Fig. 2b, 100 cells were randomly subsampled from each cell type to generate the heatmap. [[Code](https://github.com/pachterlab/CWGFLHGCCHAP_2021/blob/master/notebooks/Preprocessing/cDNAFiltering/kallistoBusRuns_StarvAndStim.ipynb/)]

The stimulation cDNA data was processed with the same kallisto-bustools workflow and commands as the starvation experiment data [[Code](https://github.com/pachterlab/CWGFLHGCCHAP_2021/blob/master/notebooks/Preprocessing/cDNAFiltering/kallistoBusRuns_StarvAndStim.ipynb)]. We initially used Louvain clustering to also filter low UMI count clusters that were then removed from downstream analysis [[Code](https://github.com/pachterlab/CWGFLHGCCHAP_2021/blob/master/notebooks/StimulationAnalysis/stimulation_Analysis.ipynb)], [[Code](https://github.com/pachterlab/CWGFLHGCCHAP_2021/blob/master/notebooks/StimulationAnalysis/deSeq2Analysis_StimulationResponse.ipynb)].

### Distance-based Comparative Analysis of Clusters [Code]

*We first used L*_*1*_ distances between starved and control cells (within each cell type) to assess how comprehensive our cell type designations were (Fig. 4). Centroids for a given cell type were calculated for starved and control cells separately, in PCA-reduced space (60PC coordinates for each cell as opposed to the raw gene expression matrix). The centroid vectors are represented as *c*_*s*_ and *c*_*c*_ for the Starved and Control cells respectively. The *L*_*1*_ distance (*d*) between them was calculated as the sum of absolute difference between the centroid coordinates:

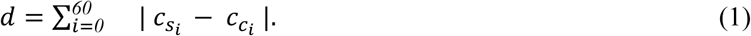

These intra-cluster distances were then compared to the pairwise *L*_*1*_ distances between the cell types, with centroids calculated for all cells in a given type (*c*_*1*_ and *c*_*2*_ for cell types 1 and 2) in the same manner as described above. The *L*_*1*_ distances were then calculated for all possible pairs of these cell type centroids using (1). The distributions of the inter- and intra-cluster distances are shown in (Fig. 4). We chose to use the *L*_*1*_ distance metric as it tends to better retain relative distances in high dimensions, particularly in comparison to the commonly used Euclidean distance or other higher *L*-norms^64,65^.

We then used the stimulation dataset to assess the validity of the clusters/cell types generated with the starvation data. We created a joint representation of the two datasets by using a concatenated cell-by-gene matrix including only the genes highly variable in both the control and starved datasets,, and used 70% of the starvation data to train a k-nearest neighbor (KNN) classifier (using sklearn’s KNNClassifier with *k* = 15) to assign cluster labels to the remaining starvation cells and to the stimulation dataset. This showed that the stimulation cells’ labels from their neighbors in starvation data were assigned at the same accuracy as the test starvation data, meaning that clusters from the starvation data are applicable to the stimulation dataset and capture the main features of the stimulation dataset to the same extent (Supplementary Fig. 5a).

We also examined batch effects in the stimulation experiment. As with the starvation experiment, the *L*_*1*_ metric was utilized to visualize the magnitude of batch effect within the multiplexed experiments compared to between experiments (Supplementary Fig. 10). We used the merged representation (used for the KNN assignment) between both experiments to find the average pairwise distances between cell types of control condition individuals within the starvation experiment, within the stimulation experiment, and across both experiments. We found that cell type distances between organisms were reduced within multiplexed experiments compared to distances across experiments.

We also used the merged atlas for determination of strong *in situ* markers, mainly for gastrodermal cell types, since highly related subtypes share many marker genes.

### RNA-seq Analysis & Clustering with MARIMBA Annotation

To make our dataset more easily searchable with the MARIMBA v.1 (created on 05/30/2016) transcriptome annotation^21^, we generated single-cell gene count matrices with respect to transcript sequences distributed via the MARIMBA website ([link](https://data.caltech.edu/records/1830)). The gene count matrices were generated using the same kallisto-bustools workflow, and compared the application of the previous clustering/cell type assignment (including the overlap in differentially expressed genes delineating these clusters) to validate the quantification derived from the Trinity/Cuffcompare assembled transcriptome (Supplementary Fig. 18). [[Code](https://github.com/pachterlab/CWGFLHGCCHAP_2021/blob/master/notebooks/SearchAndPlotInteractive/cufflinksAnnosAnalysis.ipynb)]

We also produced a notebook for visualization of gene expression in this dataset to facilitate its use in future studies. This establishes a code base for rapidly and transparently processing and comparing single-cell datasets with future transcriptome annotations. [[Code](https://github.com/pachterlab/CWGFLHGCCHAP_2021/blob/master/notebooks/SearchAndPlotInteractive/exploreCufflinksData.ipynb)]

### Neural Analysis [Code]

We clustered all cells within the broad class of ‘Neural’ with Louvain clustering to obtain distinct subpopulations (labeled in Fig. 2a). Markers were determined with Scanpy’s rank_genes_groups function (using the Wilcoxon test) for each subpopulation (Supplementary Table 3).

Marker genes from neural clusters, with predicted signal peptides^66^, were screened for candidate neuropeptide cleavages sites (regular expression G[KR][KRED]). In cases where a sequence had more than one match to this motif, the 6 residues immediately N-terminal to the motif were inspected for similarity to each other -when similarity was present, the protein was considered a neuropeptide candidate. Predicted sequences can be found in Supplementary Table 3.

### Pseudotime Analysis [Code]

We selected cells from cell types of interest (stem cells, nematocytes, and neuronal cells) and used diffusion mapping^67^ to create a reduced dimension representation of cells, along with Scanpy’s dpt function which uses geodesic distance along the graph of cells (in the determined ‘diffusion component’ space). We then computed a PAGA-based embedding to visualize the cells in the context of the different trajectories with a ForceAtlas2 layout. To determine which genes constituted the important features in a given pseudotime trajectory, we implemented a method based on the random forest method used in the dynverse R package for extracting ‘important’ genes^68^. A random forest regression model implemented with sklearn’s random_forest_regressor was used to identify genes that were good predictors of the generated pseudotime values (grouped into quantiles). This was run for each of the two inferred trajectories separately (stem cells to nematocytes, and stem cells to neurons). The training set consisted of 80% of the genes’ expression data, within which 80% was used for optimizing the model, and 20% was used to evaluate the mean squared error (MSE). Both trajectory models had an *R*^*2*^ of 0.85 or greater. The remaining 20% of data from the full dataset was used to calculate the gene-wise permutation importance scores, providing a ranking of each feature (gene) in terms of its contribution to the model’s predictive capabilities using sklearn’s permutation_importance. In the ranking, positive scores indicate importance, with the importance *i*_*j*_ for gene *j* being the difference between the original score s (MSE) calculated from the original model, and the average score across *K=5* random permutations of the feature columns in the validation dataset:

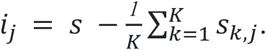

Genes with non-zero (positive) scores in the permutation test were retained and ranked (Supplementary Table 3).

### Perturbation Response Analysis

#### Extracting DE (perturbed) genes

We used a likelihood ratio test (LRT) with negative-binomial based models of each gene’s expression in DESeq2^69,70^, with the reduced model not including the condition label (fed or starved) and single-cells treated as individual replicates. There are other approaches that can be used^71,72^ ; we found that this method yielded many biologically relevant genes, which we could validate via the *in situ* experiments. All nonzero-expression genes were used for analysis. From the LRT we obtained *p*-values (corrected with the Benjamini-Hochberg multiple testing correction method across the number of genes tested) to determine if the gene’s expression was significantly affected by the condition, and thus the perturbation. Genes with alpha < 0.05 and |log2FC| > 1 were selected as significant. We used the parameters sfType=“poscounts”, minmu=1e-6, minReplicatesForReplace=Inf in the DESeq2 model. Clusters with greater than 100 cells in each condition were subsampled evenly from fed and starved cells, to reduce the effect of uneven cluster sizes on differential expression analysis (Supplementary Fig. 3). Clusters with less than 10 cells in any condition were not used for this analysis. To create the UpSet Plot^73^ in Fig. 2b, *p*-values from the LRT analysis were corrected for multiple testing (Bonferroni correction with n = number of clusters, since the test is assessing the intersection of genes across all cell types).[[Code](https://github.com/pachterlab/CWGFLHGCCHAP_2021/blob/master/notebooks/CellAtlasAnalysis/deSeq2Analysis_StarvationResponse.ipynb)]

#### De novo perturbed gene clusters

To cluster genes affected by perturbation (Supplementary Table 5) and obtain information on their co-expression and possible functional similarity, we transposed the cell-by-gene expression matrix (obtaining a gene-by-cell matrix) for only the aggregated perturbed genes with padj > 0.05 (adjusted across genes and clusters) from the DeSeq2 analysis. We used Louvain clustering on the genes expression matrix, identifying both co-expressed genes and cell type specific genes, similar to Monocle’s procedure for detecting gene-modules^53^. We then used the aggregated information from the ‘modules’ to determine putative functions/response types. The topGO weight algorithm^74^ was used to determine gene ontology (GO) terms that were significantly enriched in each gene module compared to GO terms in all other groups, with significance threshold alpha <0.05. The *p*-values were also adjusted for multiple testing over the number of different gene modules using Bonferroni correction, and only significant GO terms were used to label the response types among the modules (Supplementary Fig. 16; Supplementary Table 5). [[Code](https://github.com/pachterlab/CWGFLHGCCHAP_2021/blob/master/notebooks/CellAtlasAnalysis/starvation_Analysis.ipynb)]

## Data Availability

All raw sequencing and processed data files used for analysis are available from CaltechData with links provided via the notebooks in the code repository.

## Software used

[Cell Ranger 3.0.1](https://support.10xgenomics.com/single-cell-gene-expression/software/overview/welcome)

[Trinity-v2.8.4](https://github.com/trinityrnaseq/trinityrnaseq/releases)

[Cufflinks v2.2.1](http://cole-trapnell-lab.github.io/cufflinks/)

[kallisto v0.46.2](https://pachterlab.github.io/kallisto/)

[bustools v0.40.0](https://www.kallistobus.tools/)

[anndata 0.7.5](https://pypi.org/project/anndata/)

[louvain 0.7.0](https://pypi.org/project/louvain/)

rpy2 3.4.2

[scanpy 1.6.0](https://pypi.org/project/scanpy/)

[biopython 1.78](https://pypi.org/project/biopython/)

[pysam 0.16.0.1](https://pypi.org/project/pysam/)

[fuzzywuzzy 0.18.0](https://pypi.org/project/fuzzywuzzy/)

[numpy 0.19.5](https://pypi.org/project/numpy/)

[pandas 1.1.5](https://pypi.org/project/pandas/)

[matplotlib 3.2.2](https://pypi.org/project/matplotlib/)

[sklearn 0.0](https://pypi.org/project/sklearn/)

[scipy 1.4.1](https://pypi.org/project/scipy/)

[seaborn 0.11.1](https://pypi.org/project/seaborn/)

[requests 2.23.0](https://pypi.org/project/requests/)

[tqdm 4.41.1](https://pypi.org/project/tqdm/)

[multiprocess 0.70.11.1](https://pypi.org/project/multiprocess/)

DESeq2 1.3.0

[topGO 2.42.0](https://bioconductor.org/packages/release/bioc/html/topGO.html)

[UpSet 1.4.0](https://cran.r-project.org/web/packages/UpSetR/index.html)

## Code Availability

All code is available at https://github.com/pachterlab/CWGFLHGCCHAP_2021. The repository contains Google Colab notebooks that generate all the analyses and figures in the paper. The notebooks, which include the complete pre-processing of the raw data, provide a transparent implementation of the methods, and can be run for free in the Google cloud.

## Acknowledgements

We thank Xiaolin Da and Xiao Wang for their technical assistance, Tsuyoshi Momose for his assistance with the single-cell experimentation, the Caltech Single-Cell Profiling and Engineering Center for use of their single-cell and sequencing tools, and the Caltech Bioinformatics Resource Center for transcriptome assembly and annotation analysis. We thank A. Sina Booeshaghi for help with kallisto, bustools, the kITE demultiplexing of the ClickTag reads, and for rescuing the stimulation experiment sequencing data. We thank Jocelyn Malamy for helping to establish *Clytia* work at Caltech. We thank Sophie Peron for initial characterization of some of the cell type marker genes, Pascal Lapébie for identification of novel neuropeptide sequences, and Muriel Jager for valuable advice on the *in situ* protocol.

J.G., M.H. and L.P. were supported in part by a seed grant from the Chen Institute at the California Institute of Technology. T.C., J.G. and L.P. were supported in part by NIH U19MH114830 and NIH RF1AG062324A. We thank the Marine Resources Centre (CRBM and PIV imaging platform) of Institut de la Mer de Villefranche (IMEV), supported by EMBRC-France. The French state funds of EMBRC-France are managed by the ANR within the investments of the Future program. L.L. was supported by the Agence Nationale de la Recherche (ANR-19-CE13-0003) A.F., R.R.C, and E.H., were supported by the H2020 / Marie Skłodowska-Curie ITN “EvoCell” Grant agreement N° 766053. B.W. was supported in part by a Howard Hughes Medical Institute Fellowship of the Life Sciences Research Foundation and in part by NIH K99NS119749. This work was in part supported by the Whitman Center of the Marine Biological Laboratory in Woods Hole, MA, and a visiting grant from EMBRC-France. D.J.A. is an Investigator of the Howard Hughes Medical Institute.

## Author Contributions

T.C., B.W., J.G., R.R.C., E.H., D.J.A, and L.P. conceived the experiments. J.G. and M.H. developed cell dissociation, fixation, and labeling procedures compatible with the 10X Genomics platform. T.C., B.W., and J.G. performed the single-cell experiments. B.W., A.F., L.L., and S.C. performed the *in situ* hybridization and other microscopy experiments. F.G and R.R.C. performed bioinformatics analysis including assembly and annotation of the transcriptome. T.C. and J.G. wrote scripts for processing the data and code for the analysis. T.C. developed the Google Colab notebooks. T.C., B.W., J.G., A.F., L.L.,R.C., E.H., D.J.A, and L.P. analyzed and interpreted the data. T.C., B.W., J.G., A.F., L.L., R.R.C., E.H., D.J.A, and L.P. contributed to writing and editing the manuscript.

