## Supplementary Materials for "Whole Animal Multiplexed Single-Cell RNA-Seq Reveals Plasticity of *Clytia* Medusa Cell Types"

### Supplementary Material

Tara Chari[1+], Brandon Weissbourd[1,2,3+], Jase Gehring[4+], Anna Ferraioli[5], Lucas Leclère[5], Makenna Herl[6], Fan Gao[7], Sandra Chevalier[5], Richard R. Copley[5*], Evelyn Houliston[5*], David J. Anderson[1,2,3*] and Lior Pachter[1,8*]

[1] Division of Biology and Biological Engineering, California Institute of Technology, Pasadena, CA 91125

[2] TianQiao and Chrissy Chen Institute for Neuroscience, Pasadena, CA 91125

[3] Howard Hughes Medical Institute, California Institute of Technology, Pasadena, CA 91125

[4] Department of Genome Sciences, University of Washington, Seattle, Washington 98195

[5] Sorbonne Université, CNRS, Laboratoire de Biologie du Développement de Villefranche-sur-mer (LBDV), 06230 France

[6] University of New Hampshire School of Law, Concord, New Hampshire, 03301

[7] Caltech Bioinformatics Resource Center, California Institute of Technology, Pasadena, CA 91125

[8] Department of Computing and Mathematical Sciences, California Institute of Technology, Pasadena, CA 91125

+ Authors contributed equally

* Authors to whom correspondence should be addressed

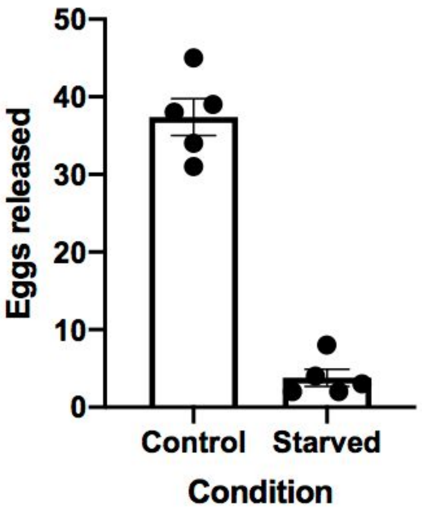


**Supplementary Figure 1:** Number of egg cells released in the daily spawning cycle from 5 control and 5 starved medusae on day four of starvation. Animals used are not the same as the individuals in the sequencing experiment, however the same protocol as in Methods was used to conduct the starvation. Error bars denote standard deviation.

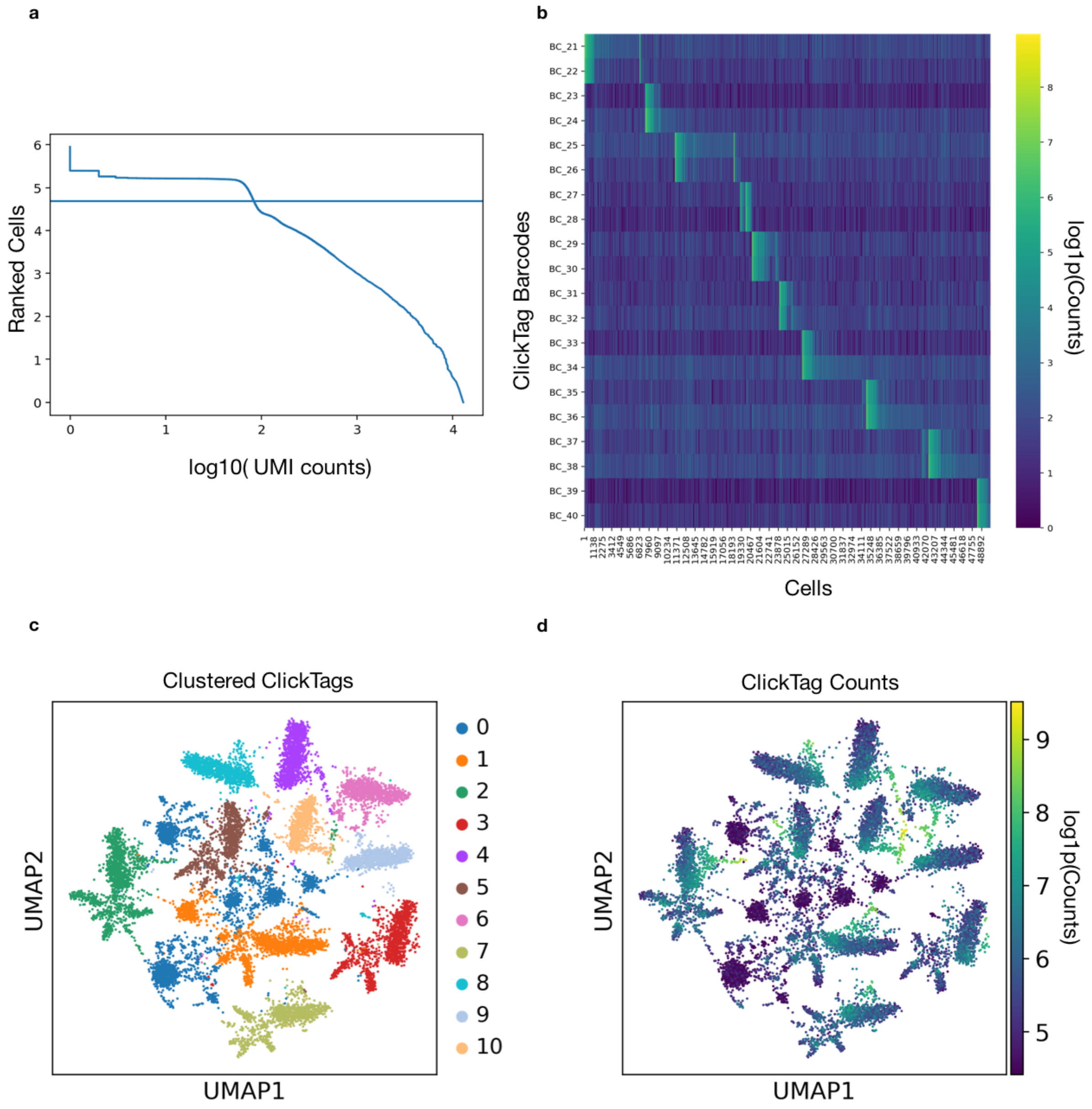


**Supplementary Figure 2: a)** Knee-plot for ranked cell barcodes (ranked according to number of UMIs) versus ClickTag UMI counts per cell barcode. Line denotes filtering/selection of the top 50,000 cells. **b)** Heatmap of counts for ClickTag barcodes associated with cell barcodes. Pairs of barcodes correspond to the ClickTags added to each organism's samples (Supplementary Table 2). **c)** Louvain clustering of ClickTags based on counts across cells, corresponding to the 10 labeled organisms. **d)** Log counts of ClickTags. High count clusters correspond to cells of the 10 dual-barcoded animals. [Code]

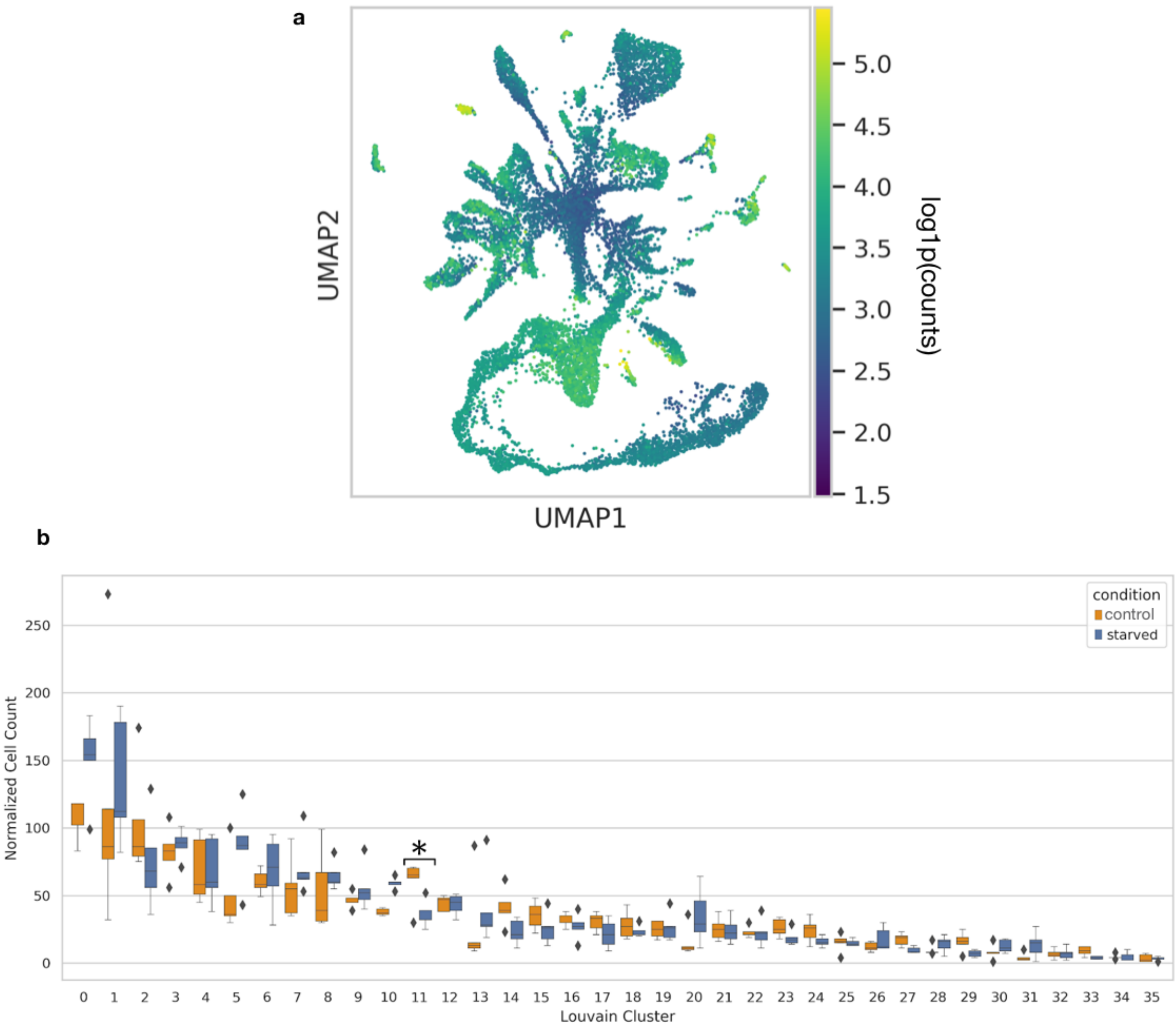


**Supplementary Figure 3: a)** UMI counts shown on log scale across cells on the UMAP embedding. [Code] **b)** One-way anova on log cell counts within each cell type (per each of the 5 individuals). Whiskers denote 1.5 IQR and diamonds denote outlier points. Pval obtained with an F test. * = $p$-value < 0.05, adjusted with Benjamini-Hochberg for FDR correction. Early nematocytes (cluster 11) show significant differences in numbers between individuals in starved vs fed conditions. [Code]

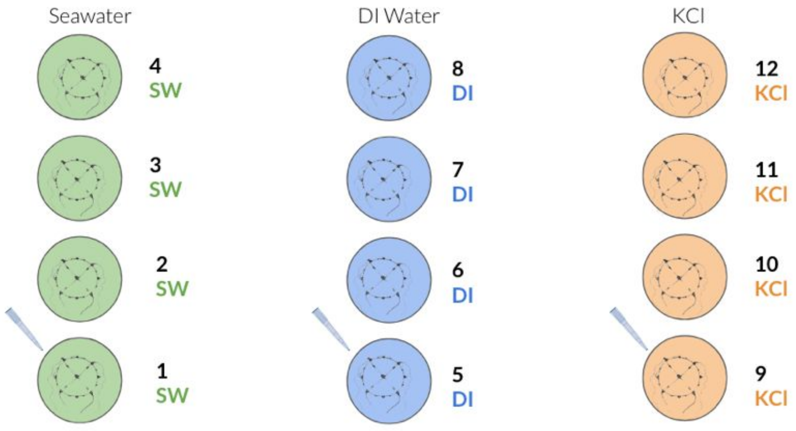


**Supplementary Figure 4:** Diagram of stimulation experiment. Four biological replicates (animals) used for each condition. SW denotes seawater (control), DI denotes deionized water addition, and KCl denotes potassium chloride addition (see Methods).

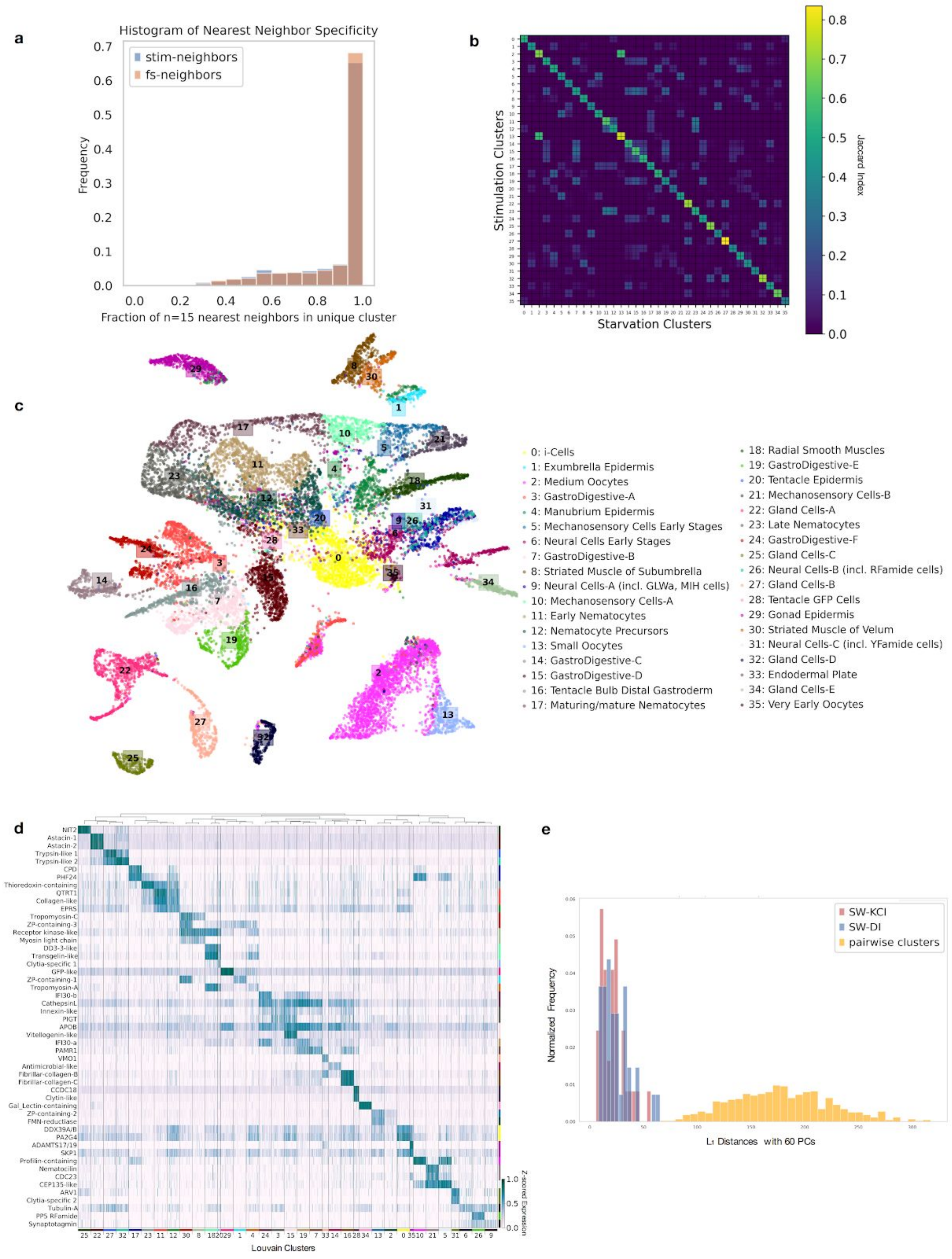


**Supplementary Figure 5: a)** Starvation data (training set) used to select nearest neighbors from left-out starvation data and all the stimulation data with the fraction of nearest neighbors selected for each dataset that were assigned to the same cell type reported (see Methods). **b)** Top 100 markers from the stimulation experiment with cluster labels applied from K-Nearest Neighbors (see Methods) compared to top 100 markers per cell type from starvation clustered data. **c)** UMAP/PAGA embedding of stimulation data with applied cell type labels. **d)** Heatmap of markers from Figure 2b shown for stimulation data. **e** Inter- and intra- cluster distances shown for control (SW) versus perturbed cells for each perturbation (see Methods). The cell type with largest internal distance (but non-overlapping) is 20 (tentacle epidermis). [Code a,e] [Code b-c]

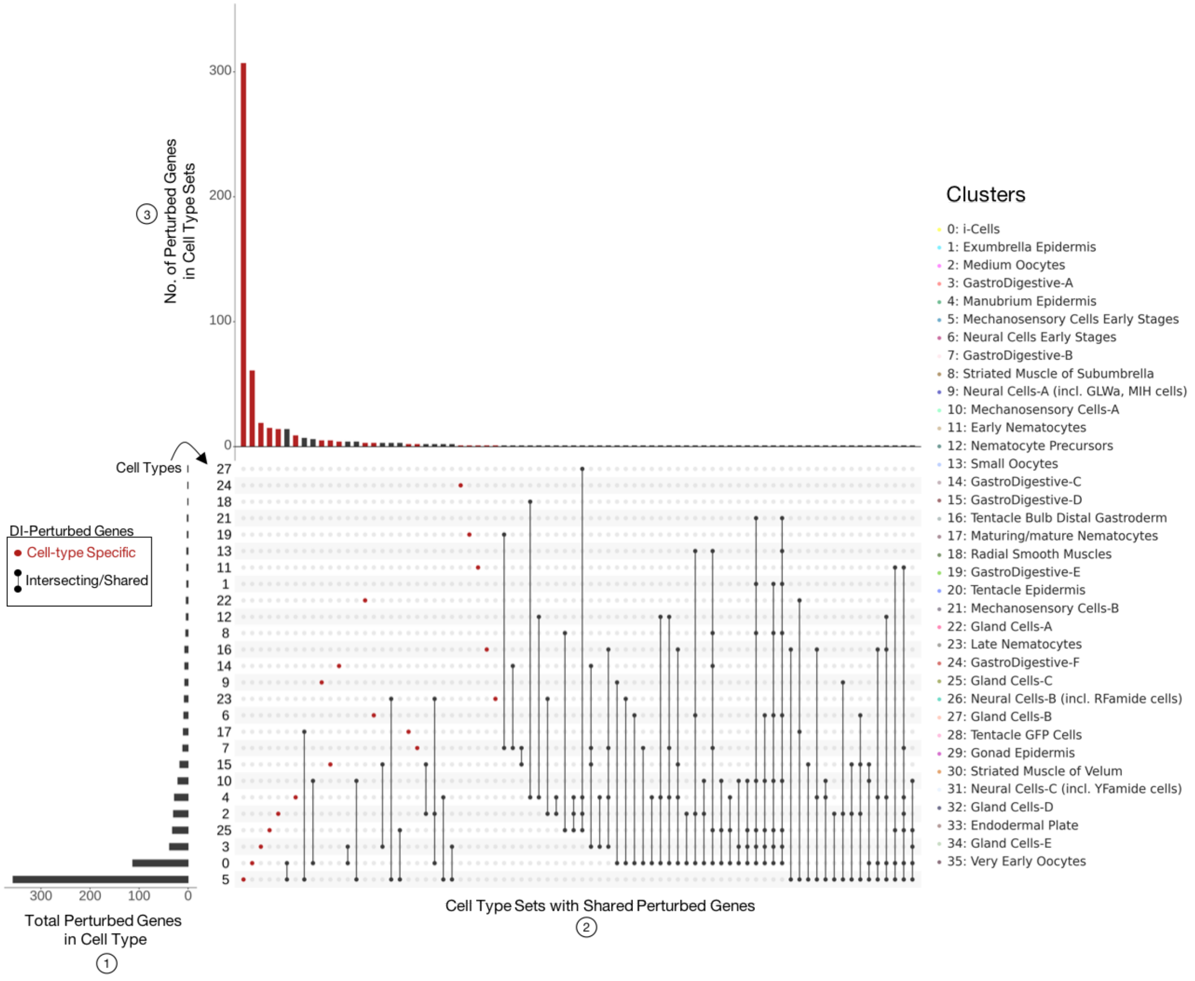


**Supplementary Figure 6:** Cell type specificity of perturbation response to DI addition from DeSeq2 analysis. UpSet plot visualization for intersecting sets of 'perturbed' genes. The barplot in 1) shows the number of genes that are differentially expressed under DI stimulation ('perturbed' genes) for each cell type. This is the cardinality or size of each cell type's set of perturbed genes. 2) The connected dots represent intersections (overlaps) between the sets of each cell type's perturbed genes. Genes with DI differential expression in only one cell type are denoted by the red, unconnected dots. 3) The top bar plot is the number of perturbed genes within these sets of shared perturbed genes, i.e. the cardinality of the intersections in 2. Cell type 5 (developing mechanosensory cells) had the most (unique) perturbed genes. [Code]

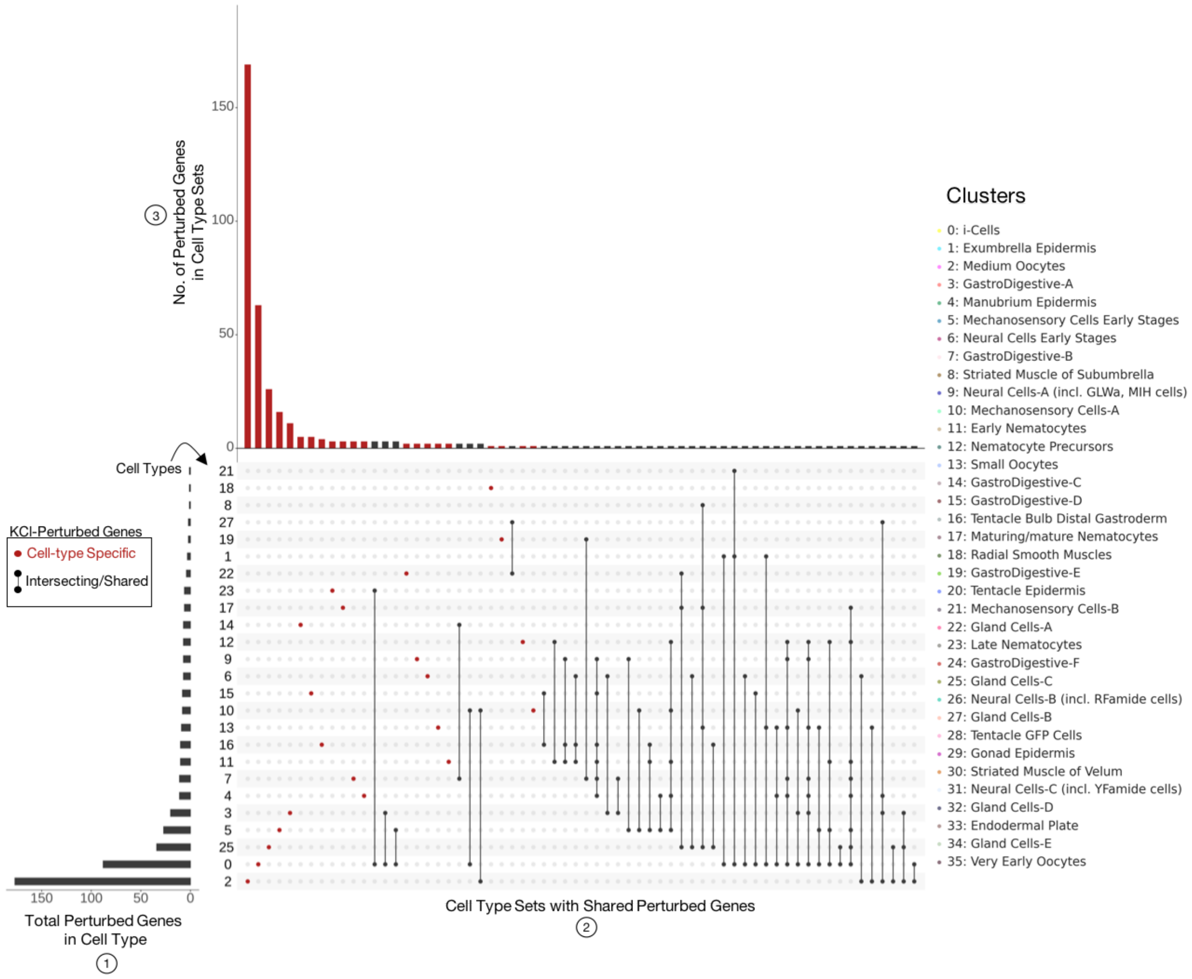


**Supplementary Figure 7:** Cell type specificity of perturbation response to KCl addition from DeSeq2 analysis. UpSet plot components are the same as in Supplementary Fig. 6. Cell type 2 (medium oocytes) had the most (unique) perturbed genes. [Code]

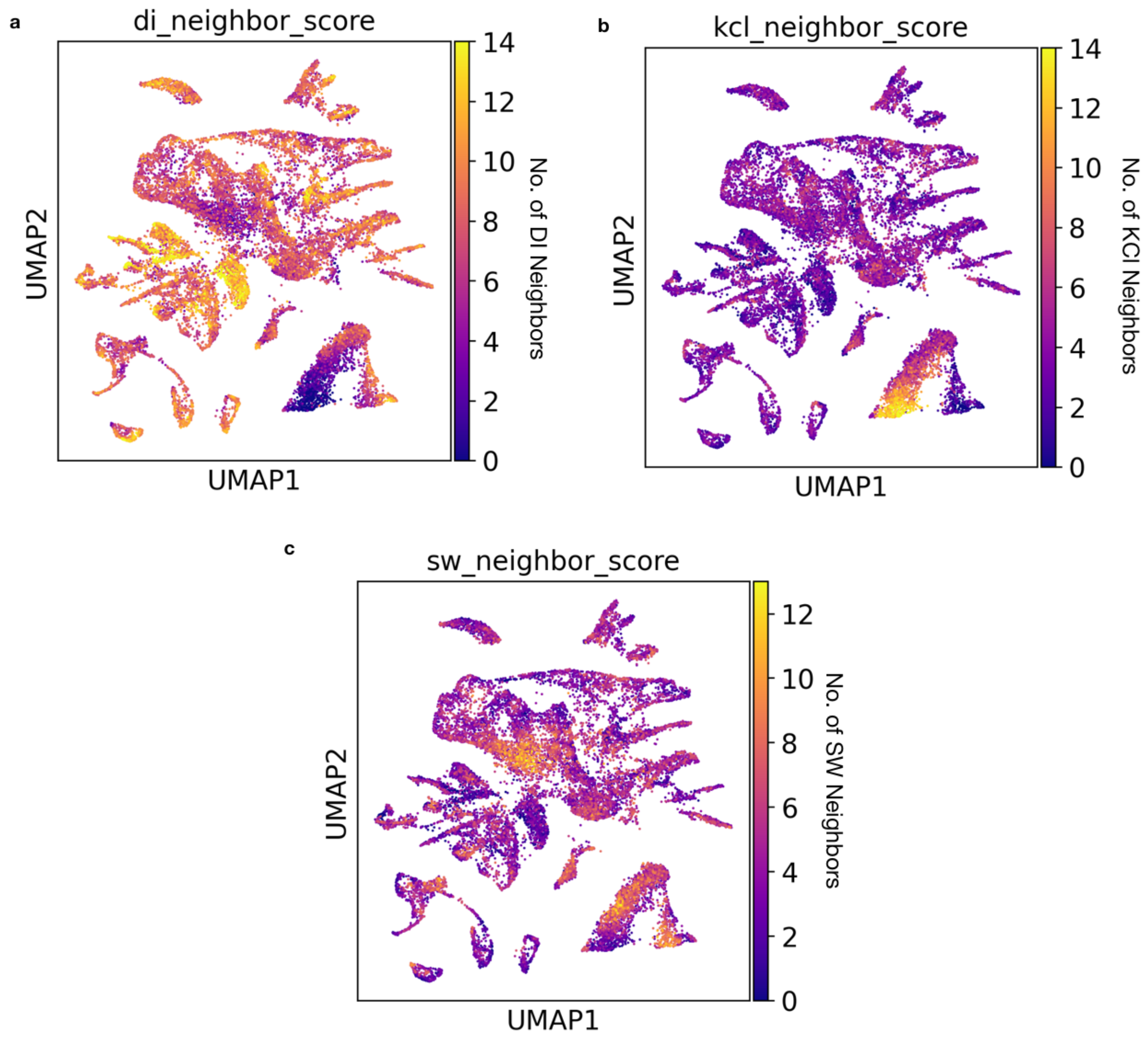


**Supplementary Figure 7: a)** Neighbor score plotted, in which cells are colored by how many of their 15 nearest neighbors are DI-treated cells. **b)** Cells colored by how many of 15 nearest neighbors are KCl-treated cells (clustered in cell type 2, medium oocytes). **c)** Cells colored by how many of 15 nearest neighbors are SW (control i.e. sea-water) cells. [Code]

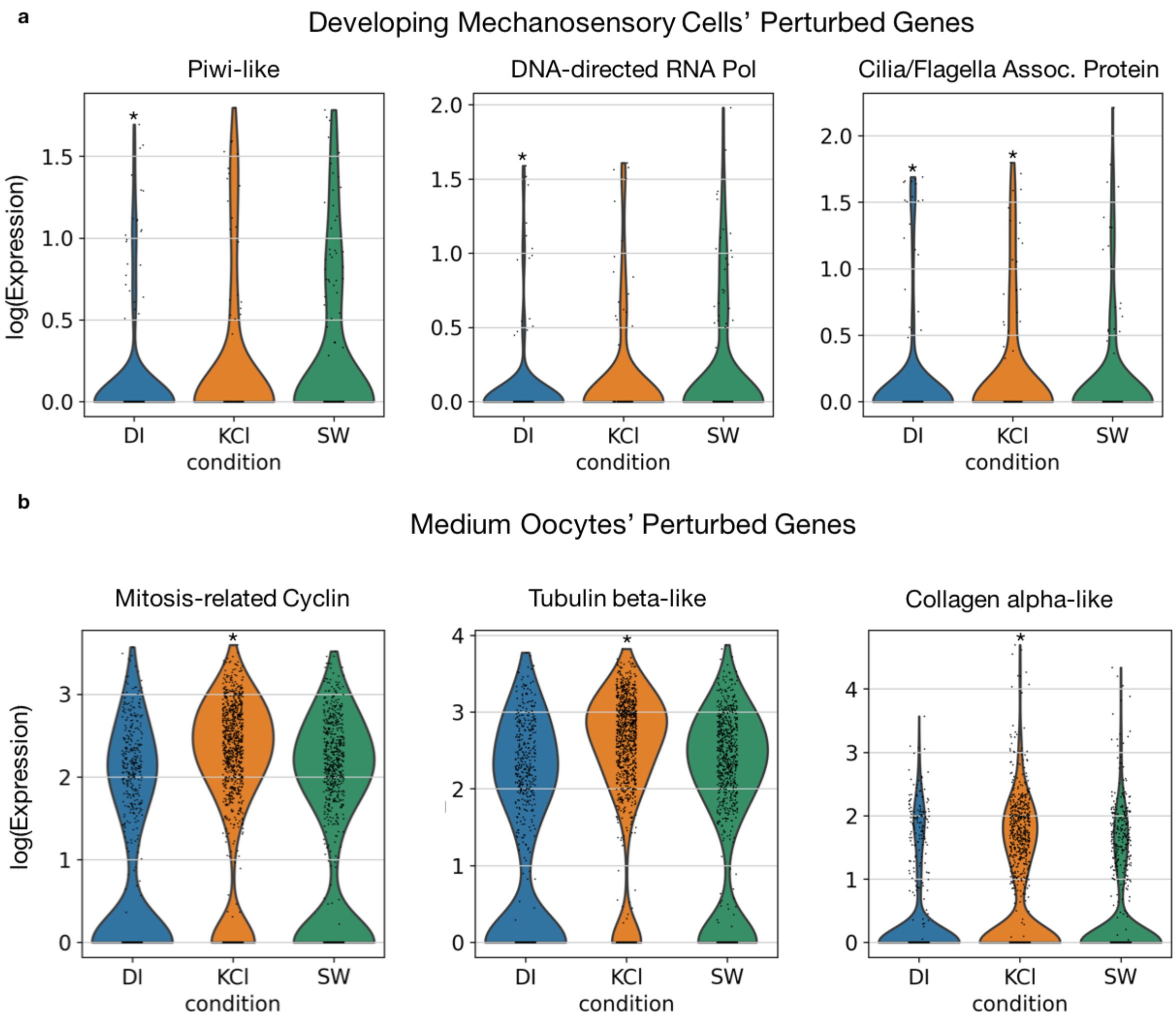


**Supplementary Figure 9: a)** Violin plots showing three examples of differentially expressed (particularly down-regulated) genes under DI perturbation in cell type 5 (Developing Mechanosensory Cells). However, there is shared down-regulation in KCl for the Cilia related gene. **b)** Violin plots showing examples of differentially expressed (up-regulated) genes under KCl stimulation in cell type 2 (medium oocytes). * indicate significant differential expression (vs the control SW), adjusted p-value < 0.05. [Code]

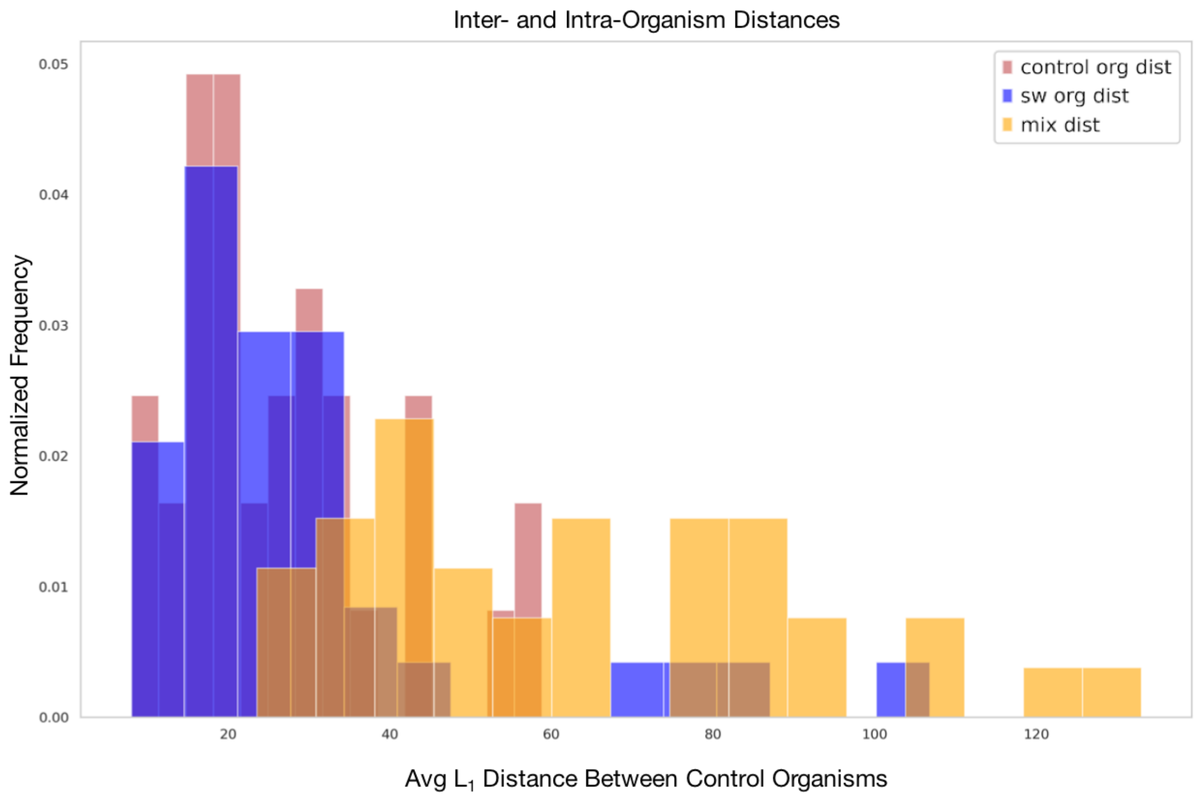


**Supplementary Figure 10:** Histograms of average pairwise $L_1$ distances between cell types of control (fed) animals within (red and blue) and across (yellow) experiments for visualization of batch effect (see Methods). [Code]

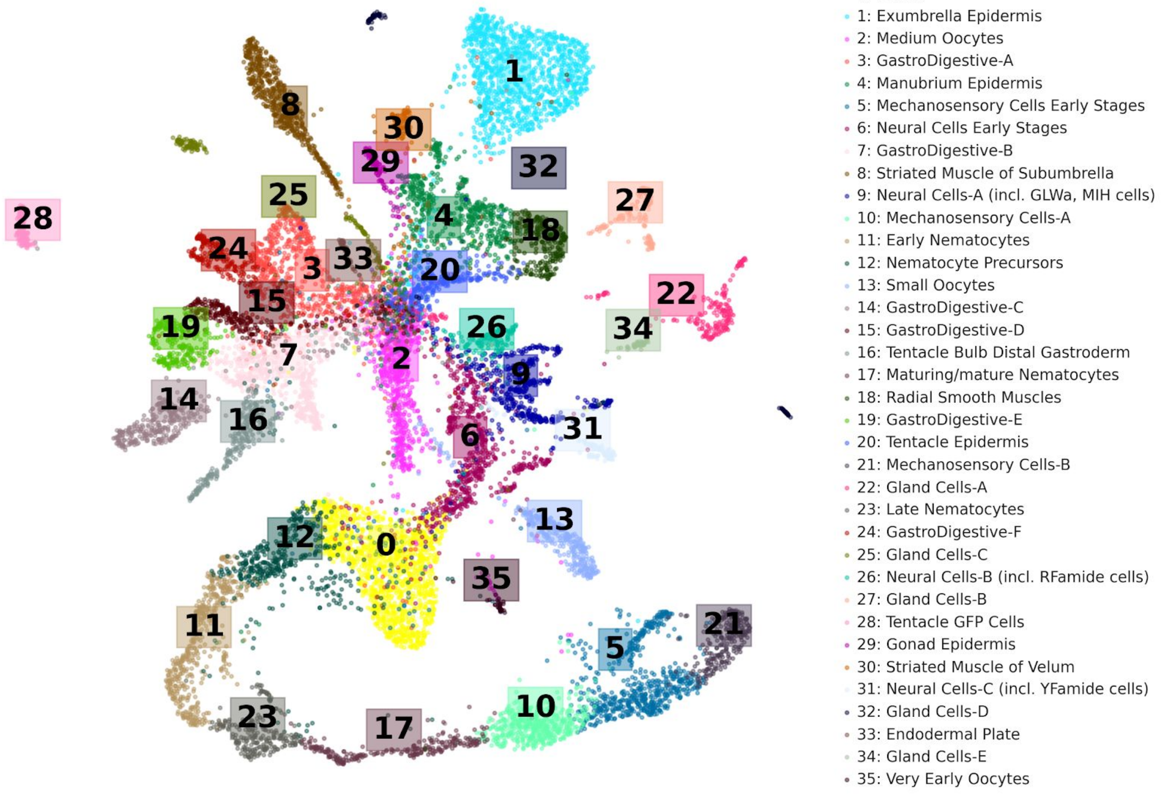


**Supplementary Figure 11:** Cell Atlas with all 36 clusters/cell types labeled with UMAP/PAGA embedding from the starvation experiment data (see Methods). [Code]

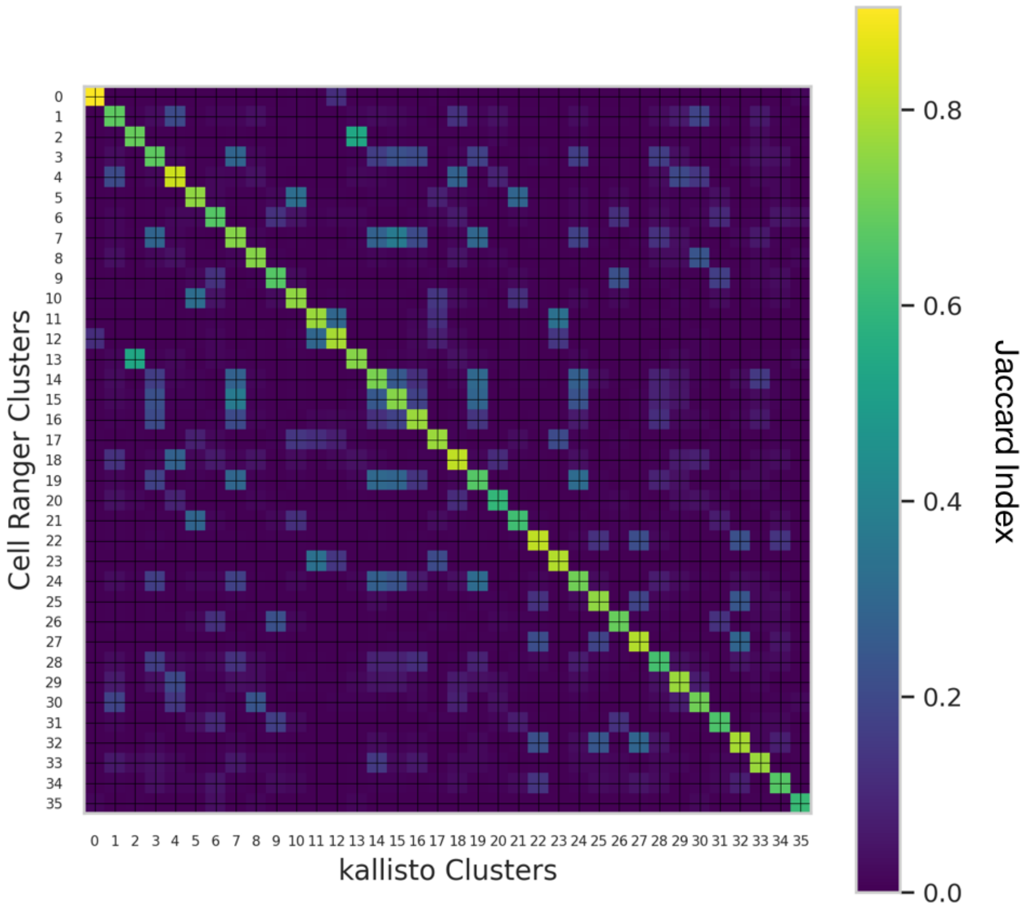


**Supplementary Figure 12:** Pairwise overlap of the top 100 marker genes from the original 36 Cell Ranger clusters applied to kallisto-bustools processed starvation data versus the top 100 markers from the original Cell Ranger clustered data. The coloring reflects the Jaccard Index of each top marker gene set (number of genes in the intersection divided by the number of genes in the union). [Code]

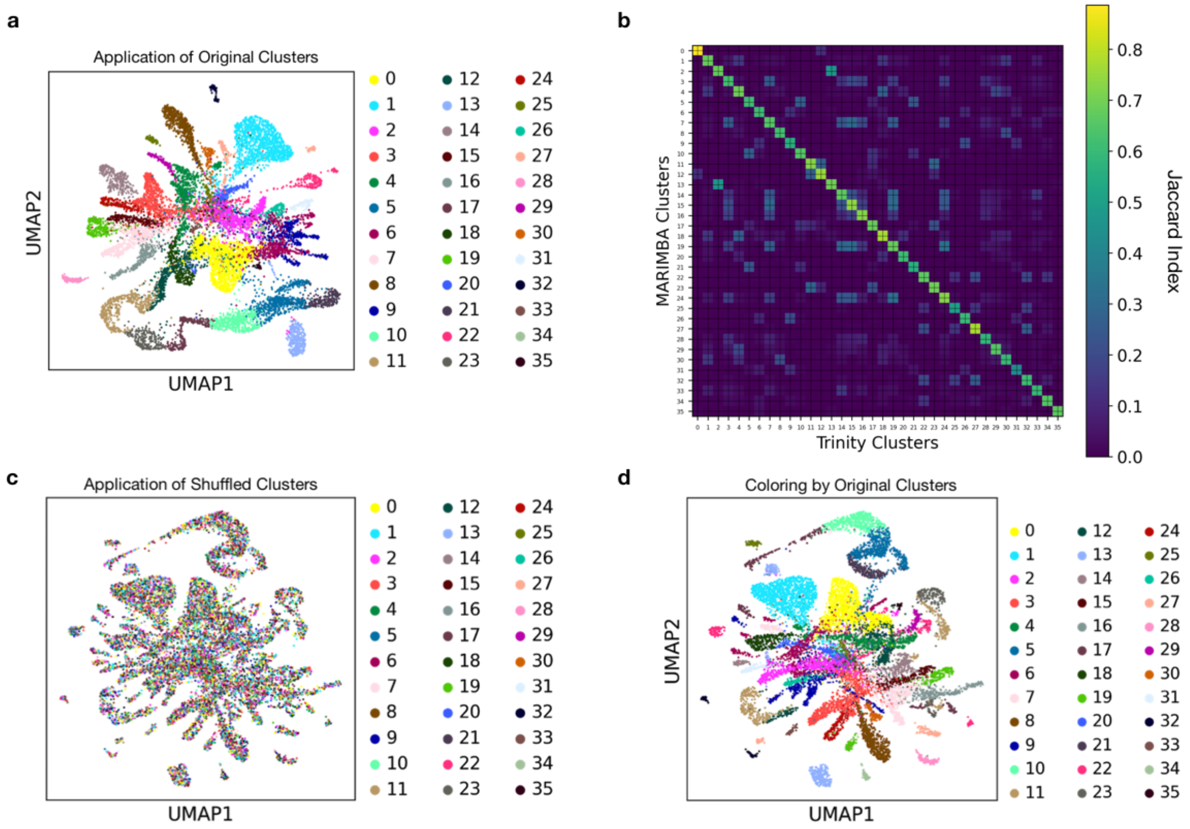


**Supplementary Figure 13: a)** UMAP/PAGA embedding of the cell-by-gene counts derived from processing the scRNA-seq data with respect to MARIMBA v.1 with the previously derived 36 cell type labels applied to the cells (see Methods). **b)** The Jaccard index for the top 100 markers from the MARIMBA v.1 annotation derived clusters versus from the Trinity/Cuffcompare annotation derived clusters. **c)** UMAP/PAGA embedding of the count matrices derived from MARIMBA v.1 annotations with scrambled cell type labels used as PAGA input. **d)** Embedding from **c** colored according to the cell types derived from Trinity/Cuffcompare annotations. [Code a-c]

| GENERAL IDENTITY | CLUSTER | KEY DIAGNOSTIC GENES | CELL TYPE |
|---|---|---|---|
| Epidermal / muscle | 1 | TPMA, GFP3[38] | Exumbrella epidermis |
| | 4 | ST MyHCa, ST MyHCb, GFP4[38] | Manubrium epidermis |
| | 8 | ST MyHCa, TPMB | Striated muscle of subumbrella |
| | 18 | TPMA | Radial smooth muscle |
| | 20 | TPMA, ST MyHCa, Wnt2[34], | Tentacle epidermis |
| | 29 | TPMA, ST MyHCb, GFP4[38] | Gonad epidermis |
| | 30 | ST MyHCa , TPMB, Peroxidase | Striated muscle of velum |
| Gastroderm | 3 | CathepsinL | GastroDigestive A |
| | 7 | CathepsinL, Nucleoside hydrolase | GastroDigestive B |
| | 14 | CathepsinL, BP10-like , Fibulin | GastroDigestive C |
| | 15 | CathepsinL, VCBS repeat protein | GastroDigestive D |
| | 19 | CathepsinL, FibCdom-2 | GastroDigestive E |
| | 24 | CathepsinL, DDAH2 | GastroDigestive F |
| | 16 | FibColl-2 | Tentacle bulb distal gastroderm |
| | 33 | FibColl-2, FibColl-1, Fibulin | Endodermal plate |
| Bioluminescent cells | 28 | GFP2b [38] | Tentacle GFP cells |
| Stem cell /Germ cells | 0 | Nanos1[33], Piwi[33], Vasa[33], Znf845 | i-cells |
| | 35 | Nanos1[33], Piwi[33], Vasa[33], Scyp1[8] | Very early oocytes |
| | 13 | Nanos1[33], Piwi[33,] Vasa[33], GFP2a[38], Clytin2[38] | Small oocytes[a] |
| | 2 | Nanos1[33], Piwi[33], Vasa[33], GFP2a[38], Clytin2[38] | Medium oocytes[a] |
| Gland cells | 22 | ShKT-TrypB | Gland cells A |
| | 27 | C3-Lipase , Tryp-like , Fibulin | Gland cells B |
| | 25 | FibCdom-1 | Gland cells C |
| | 32 | C3-Lipase | Gland cells D |
| | 34 | Chitinase | Gland cells E |
| Nematocytes | 12 | Minicollagen 3/4[35], Znf845, Mos3 | Nematocyte precursors |
| | 11 | Minicollagen 3/4[35], Mos3 | Early Nematocytes |
| | 23 | Minicollagen 3/4[35], Mos3 | Late Nematocytes |
| | 17 | Minicollagen 3/4[35] Whirlin[42,] | Maturing/mature Nematocytes [b] |
| Mechanosensory | 5 | Lynx, LOX, Harmonin[42], Whirlin[42], Sans[42] | Mechanosensory cell early stages |
| | 10 | Lynx, Harmonin[42], Whirlin[42], Sans[42] | Mechanosensory cells A |
| | 21 | Lynx, LOX, Harmonin[42], Whirlin[42] | Mechanosensory cells B |
| Neural | 6 | ELAV, PP3[c], PP19[c], | Neural cell early stages |
| | 9 | ELAV, PP11[c], Opsin9 [55] | Neural cells A (inc. GLWamide & MIH cells) |
| | 26 | ELAV, PP5[c] | Neural cells B (inc. RFamide cells) |
| | 31 | ELAV, PP9[c] | Neural cells C (inc. YFamide cells |

**Supplementary Figure 14:** Overview of cluster identities deduced from markers including the key diagnostic genes listed. These include *Clytia* genes whose expression has been previously characterised[33-35, 38, 55] and genes with clear homology to genes with clear cell type specificity in other animals ( Scyp1= XLOC_008881, ortholog to Hydra Scyp1/ Sc4wPfr_899.g22607 [8]; Harmonin/USH-IC =XLOC_003773, Whirlin=XLOC_01192, Sans=XLOC_039341[42]). For all other genes cited, *in situ* localisation profiles are shown in Fig. 2 and/or Supplementary Fig. 14.

[a] Clusters 2 and 13 have overlapping profiles including known oocyte specific genes such as GFP 2 (XLOC_004150) and ones highly expressed in published *Clytia* "Growing" and "Fully grown" oocyte transcriptomes[37] .

[b] Cluster 17 comprises two subpopulations preferentially expressing either Minicollagen3/4 (XLOC_004102) or mechanosensory structural genes including Whirlin (XLOC_0119), likely representing maturing and mature nematocytes respectively. Minicollagen expression downregulates during nematocyte maturation[34,36] .

[c] Details of neuropeptide precursors in Supplementary Table 4.

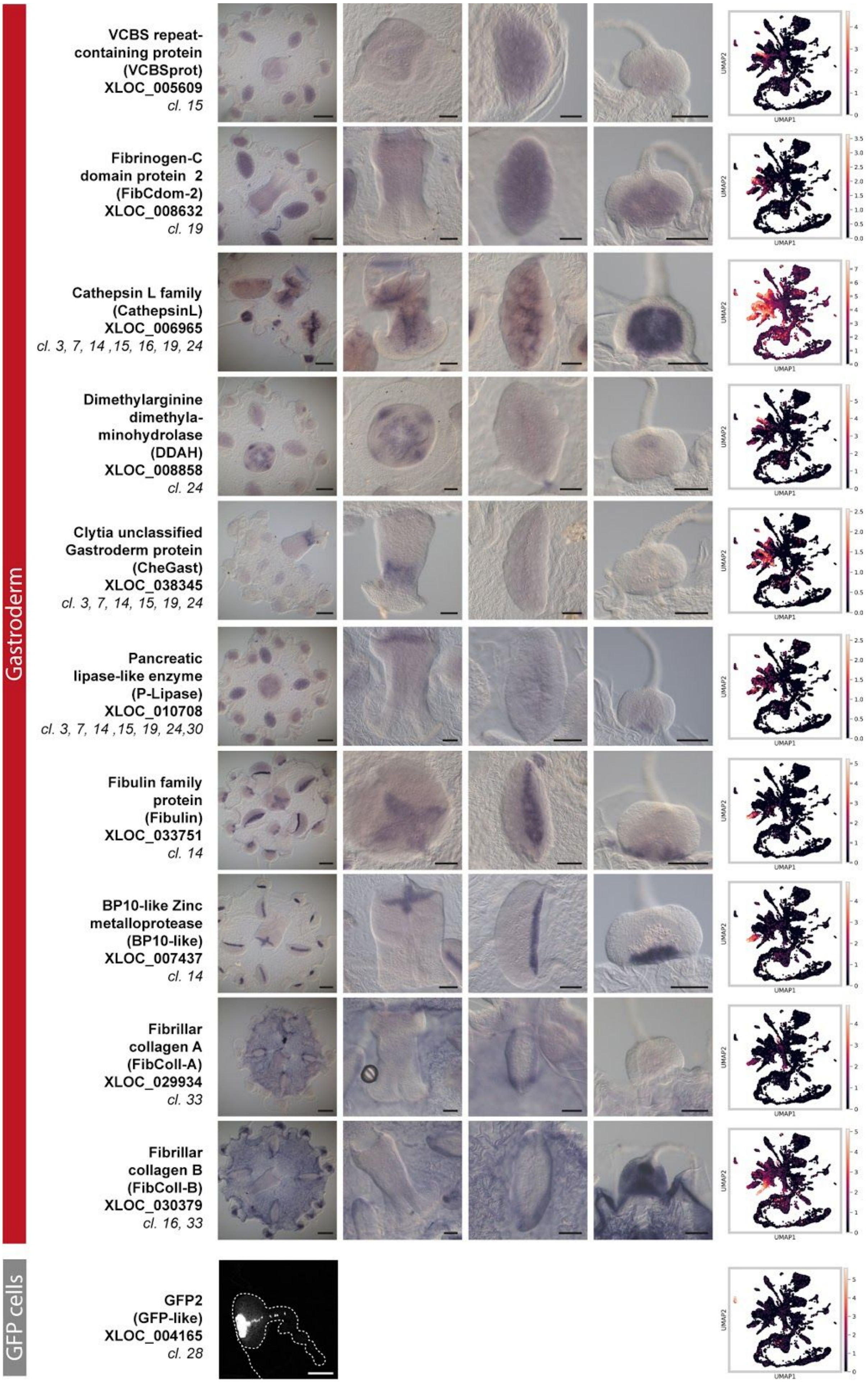


**Supplementary Figure 15:** Continued.

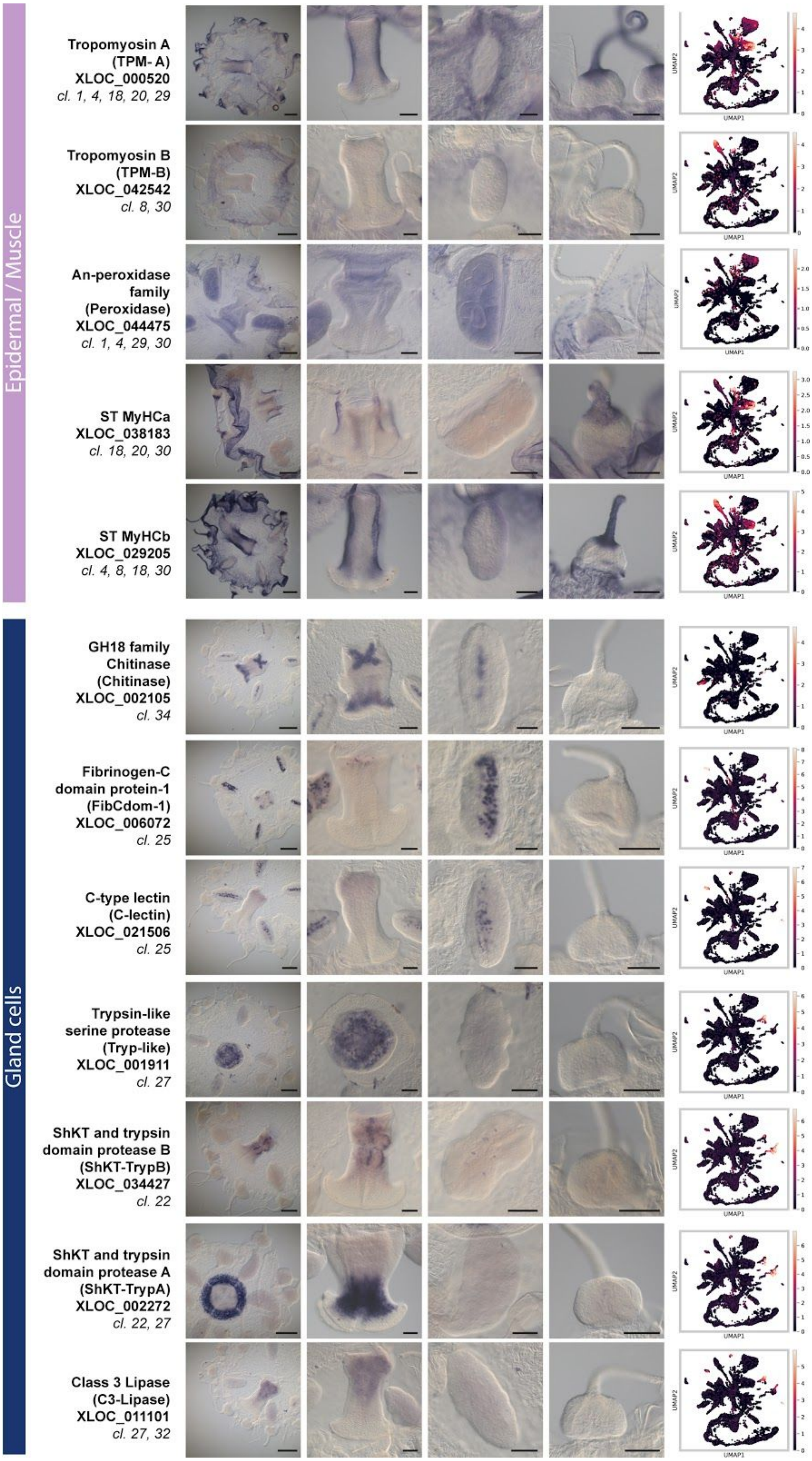


**Supplementary Figure 15:** Continued.

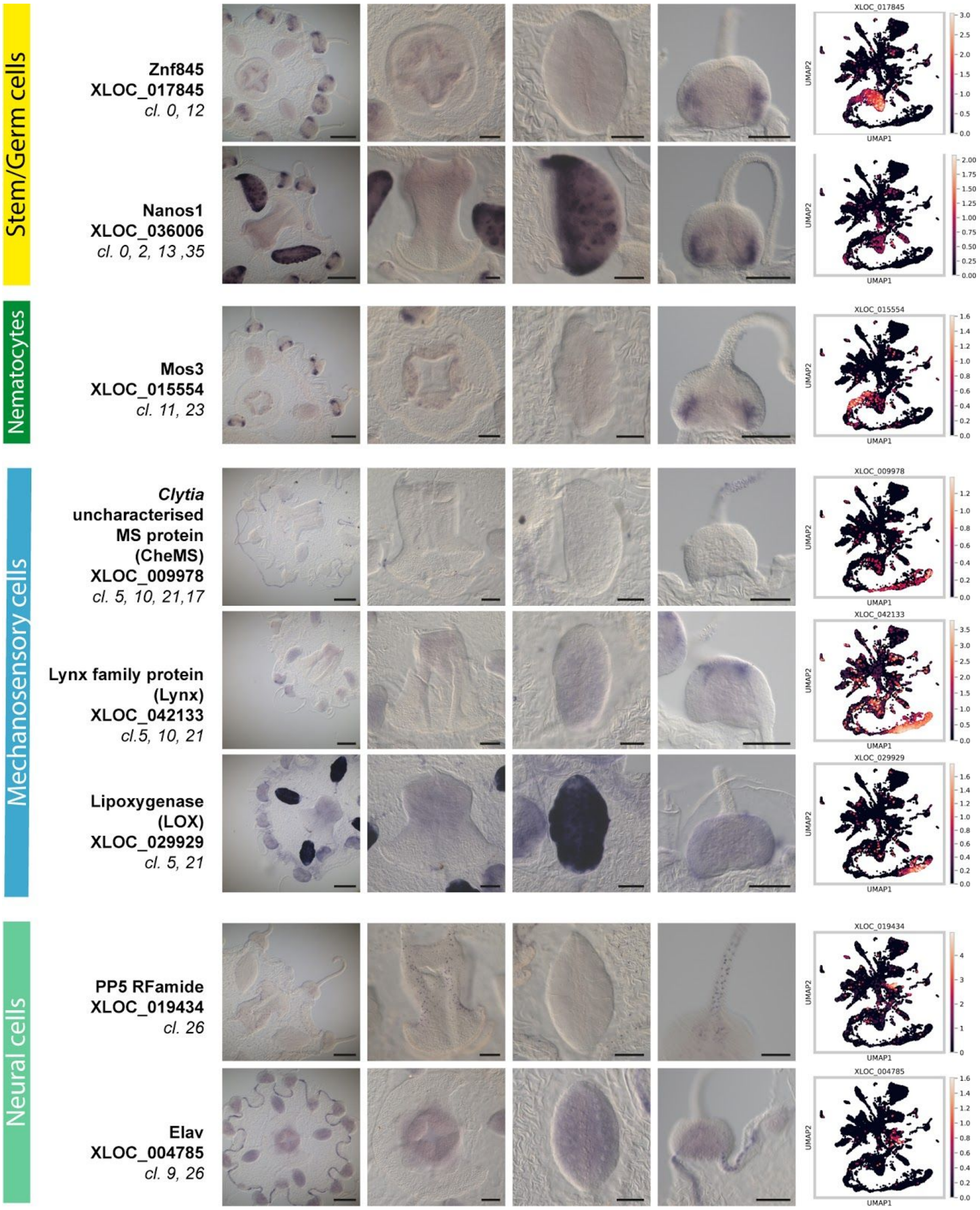


**Supplementary Figure 15:** *In situ* hybridization patterns for a selection of cluster marker genes providing spatial location of expression on the animal. Cluster IDs (cl.) indicated beneath the gene names/XLOC identifiers were assigned directly using the marker gene lists and/or by comparing gene expression profiles in the merged-experiment dataset (see Methods) . Images from left to right: whole medusa, manubrium, gonad, tentacle bulb. Right column: Gene expression for each of the marker genes represented in red in the single-cell atlas. Scale bars in the whole medusa images represent 200 µm; in the manubrium, gonad and tentacle bulb images they represent 100 µm. [Code]

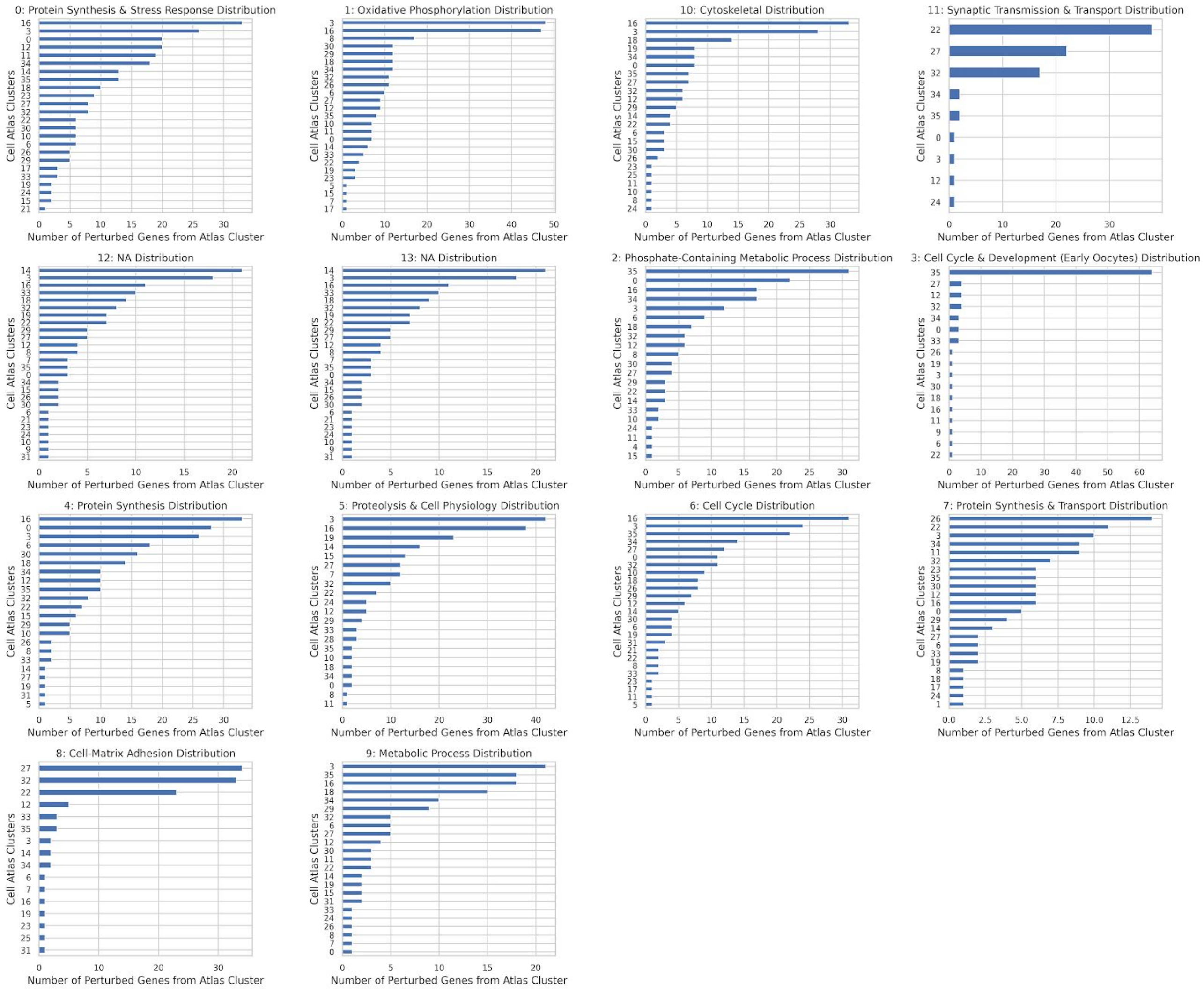


**Supplementary Figure 16:** The number of genes contributed by each of the 36 cell types to each of the gene modules.. Names for gene modules were assigned based on gene ontology (GO) term enrichment with topGO (see Methods). 'NA' is assigned to gene modules with no significant GO terms. [Code]

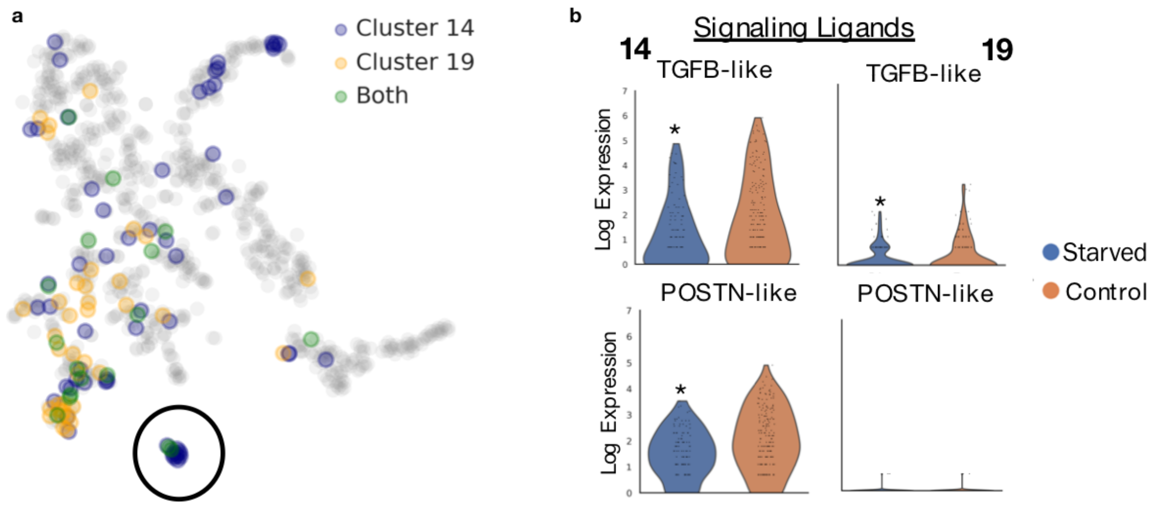


**Supplementary Figure 17: a)** Example of a gene module (module 13 from Supplementary Fig. 16, circled) with shared and unique perturbed genes between 'internally-distant' GD cell types 14 and 19. **b)** Violin plots show expression of signaling ligand related genes in this module, with shared and divergent expression between the cell types. Both cell types show downregulation of the same TGFB-like gene under starvation, however downregulation of a POSTN-like gene (related to cell adhesion and migration) is only present in 14, with low/no expression in 19 for either condition. * denote $p$-value < 0.05 for differential expression, using the non-parametric, Wilcoxon test. [Code]

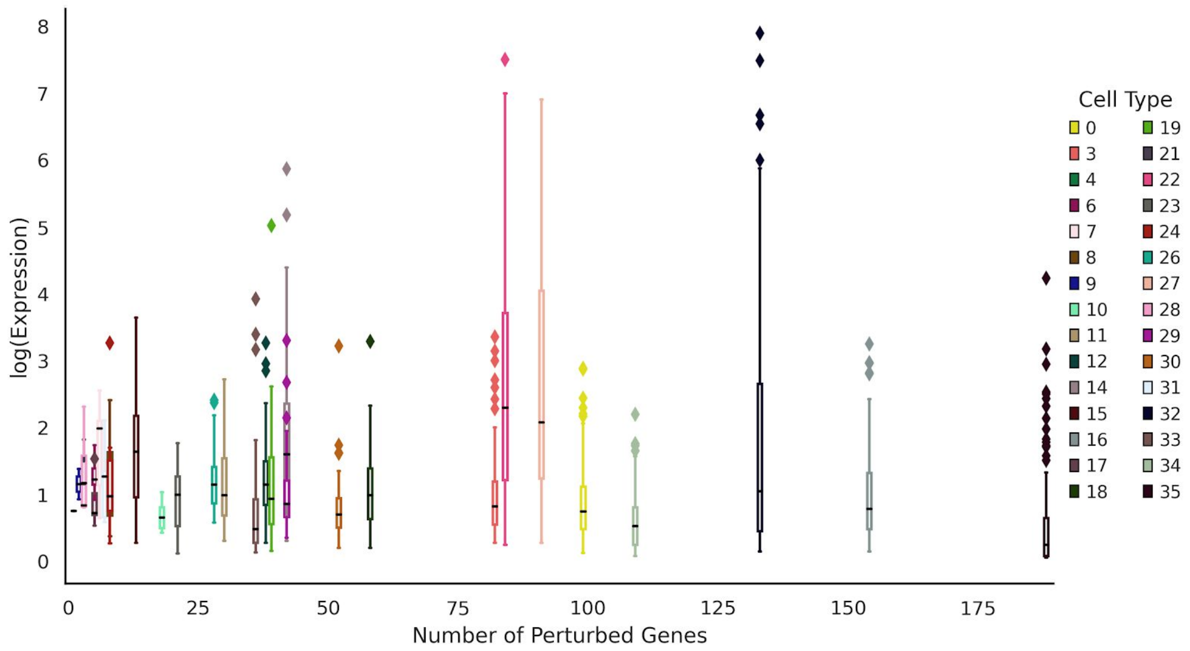


**Supplementary Figure 18:** Box-and-whisker plot showing expression level of perturbed genes (distribution across cells; mean indicated with horizontal bar) within each cell type versus the number of significantly perturbed genes found for each cell type. Whiskers denote 1.5 IQR and diamonds denote outliers. [Code]

| Experiment | No. of conditions | Number of animals | Number of high quality cells | Total Sequenced Reads | Total cost | Unique reads in high quality cells | Median UMIs per ell | Median genes detected per ell | Technology used |
|---|---|---|---|---|---|---|---|---|---|
| Starvation | 2 | 10 | 13,673 | 464,778,632 | ~$12,000 | 67,374,740 | 1,802 | 676 | 10X Chromium V2.0<br>HiSeq 4000<br>MiSeq v3 |
| Stimulation | 3 | 12 | 18,921 | 676,754,269 | ~$12,000 | 170,722,140 | 4,297 | 1,303 | 10X Chromium V3.0<br>HiSeq 4000<br>MiSeq v3 |

**Supplementary Table 1:** Multiplexed experiment(s) overview.

| Condition | Control (Fed) | Control | Control | Control | Control | Starved | Starved | Starved | Starved | Starved |
|---|---|---|---|---|---|---|---|---|---|---|
| Animal | Org 1 | Org 2 | Org 3 | Org 4 | Org 5 | Org 6 | Org 7 | Org 8 | Org 9 | Org 10 |
| Individual ClickTag | 21,22 | 23,24 | 25,26 | 27,28 | 29,30 | 31,32 | 33,34 | 35,36 | 37,38 | 39,40 |

**Supplementary Table 2:** Starvation experiment ClickTag assignments.

**Supplementary Table 3:** Designed and Predicted Sequences. All designed and predicted sequences i.e. ClickTag barcodes, *in situ* primers (with EST sequences where applicable) and predicted neuropeptide sequences.

| Condition | SW | SW | SW | SW | DI | DI | DI | DI | KCl | KCl | KCl | KCl |
|---|---|---|---|---|---|---|---|---|---|---|---|---|
| Animal | Org 1 | Org 2 | Org 3 | Org 4 | Org 5 | Org 6 | Org 7 | Org 8 | Org 9 | Org 10 | Org 11 | Org 12 |
| Individual ClickTag | 25 | 26 | 27 | 28 | 29 | 30 | 31 | 32 | 33 | 34 | 35 | 36 |
| Condition ClickTag | 37 | 37 | 37 | 37 | 38 | 38 | 38 | 38 | 39 | 39 | 39 | 39 |

**Supplementary Table 4:** Stimulation ClickTag assignments.

**Supplementary Table 5:** All marker and differentially expressed (DE) genes. Contains genes marking cell types, DE genes between starved and control cells, neuron subpopulation markers, pseudotime markers, and DE genes between stimulation conditions with functional annotations where possible.

### Supplementary Methods

#### *Full Protocol for ClickTag Sequencing*

##### *Starvation Single-cell Suspensions*

- 1x PBS at 350 mM NaCl (hypertonic PBS solution)
- MeOH (stored at -80)
- LoBind Tubes (1.5 mL)

1. In six-well plates, transfer animals sequentially to three 4 C wash baths each with 25 mL filtered hypertonic PBS (1x PBS at 350 mM NaCl). Perform all subsequent steps at 4 C. Note: This is because seawater precipitates in methanol but hypertonic PBS does not. The washes thus both prepare the animals for fixation and remove precipitating sea salts
2. After washing animals, transfer in ~400 uL to a 1 mL glass dounce homogenizer and pestle (loose) (Wheaton, USA).
3. Homogenize animals by passaging 30 times with the dounce. First transfer samples in 200 uL aliquots to Eppendorf tubes, triturated 30 times with a P200 pipet, and transfer again to a dounce (passaging 30 times) .
4. Bring volume to ~1.2 mL, and place sample on ice
5. Process corresponding samples from the control or experimental group and centrifuge at 300 x *g* for 3 minutes at 4 C.
6. Remove supernatant, leaving ~20 uL. Resuspend cells thoroughly in this volume using a P20 pipette tip.
7. Add four volumes (~80 uL) ice-cold methanol to the sample and triturate for 30-60 seconds to reduce clumping during the early stages of MeOH fixation.
8. Place samples at -20 C and store until sample labeling. Note: All samples were processed in this way, using pairs of experimental and control samples, alternating which sample was processed first, and tracking all animal characteristics with a known animal/barcode combination.
9. In between sample pairs, clean and sterilize dounce/pestles with 20% bleach, rinsed thoroughly with water, and with 70% EtOH.

##### *One-Pot Sample Labeling*

1. For each sample, perform sample labeling according to Gehring et al. 2020:
   Tags distributed as in Supplementary Table 2.
   a. 6 uL sample tag 1 (20uM)
   b. 6 uL sample tag 2
   c. 4 uL 1 mM NHS-TCO
   d. Incubate 30 mins at room temperature on a rotating platform
   e. Quench by addition of Tris-HCl (to 10 mM final) and MTZ-DBCO (to 50 uM final) using 5 uL of a 200 mM Tris-HCl, 1 mM MTZ DBCO solution (DMSO, 50 mM MTZ-DBCO, 1 M Tris-HCl pH 7.5)
2. Pool samples and supplement with 10 uM "blocking oligo" to add excess DNA for quenching the reaction. Here we use the SoIvdAR primer at 100 uM stock

(Supplementary Table 3). 10 uL of resulting samples are combined with 9 uL PBS-BSA and 1 uL DAPI 400 ug/mL for counting.

3. Final combined volume of methanol-fixed cells is ~1mL. After mixing, split combined sample in two and add 500 uL PBS-BSA (0.1% BSA) to both samples.
4. Mix and spin (1000G for 5mins). Breakspeed 5
5. Remove supernatant and resuspend in 1.2 mL PBS-BSA
6. Wash again with with PBS-BSA (optional fourth wash if desired)
7. Supernatant removed, resuspend cells in 75 uL PBS-BSA and use 10 uL for counting on Countess.
8. Count fed and starved samples with a Countess automated cell counter by brightfield and DAPI fluorescence.
    a. Using the Countess, cell concentrations are determined to roughly equate the contributions from both samples. Filter suspension with 100 micron filter before counting. Note: for this experiment, cell concentrations were determined to be 200,000 cells/mL for the starved sample and 1 M cells/mL for the fed sample
    b. To proceed, equate contributions of the starved sample (~120,000-150,000 cells) to 200,000 cells (200 uL) from the fed sample.
9. Load 10X Chromium Controller with v2 chemistry (two lanes) following manufacturer's instructions
10. Separate/process sample tag libraries after SPRI size-selection step as described in Gehring et al. 2020.
11. Sequence tag libraries on 2 lanes of an Illumina MiSeq sequencer using MiSeq v3 150 cycle kits (26 × 98-base-pair reads).
12. Pool and sequence the cDNA libraries on an Illumina HiSeq 4000 using two HiSeq 3000/4000 SBS 300 cycle kits (2 × 150-base-pair reads) Gehring et al. 2020.

*Stimulation Protocol*

- 150 mM KCl --> *KCl concentration of SW approx 10mM already*
- DI water
- 1x PBS at 350 mM NaCl (hypertonic PBS solution)
- MeOH (stored at -80)
- LoBind Tubes (1.5 mL)

Prepare samples using the same protocol as for the starvation experiment with two tags for each sample: one tag unique to each individual (1-12), and one tag unique to each condition (DI, KCl, SW) (Supplementary Fig. 4; Supplementary Table 4). The "blocking" oligos in this experiment are 5uM LCDBF and 5 uM LCDBR primers (to add excess DNA to the reaction) (Supplementary Table 3).

The washing steps prior to Countess cell counting (steps 3-7) are replaced with the steps below.

1. Final combined volume of methanol-fixed cells is ~1 mL. After mixing, split combined sample in two, and add 1.4ml 3x SCC and 0.1% BSA to both samples
2. Mix and spin (1000G for 5mins). Breakspeed 5
3. Remove supernatant and resuspend in 4 ml 3x SCC and 0.1% BSA

4. Wash again with 4ml 3x SCC and 0.1% BSA
5. Supernatant removed, resuspend cells in 150 uL PBS and 0.1% BSA and use 10 uL for counting on Countess. Continue as written in step 8.

Run samples through two lanes on a 10X Chromium Controller with v3 chemistry following manufacturer's instructions. With resulting libraries, sequence on two Illumina HiSeq lanes with 3% tags in pooled mix (cDNA and ClickTags), and run a ClickTag only sample on two Illumina MiSeq lanes.
